# Higher-order Architecture Shapes Concerted Evolution in a Y-linked repeat array

**DOI:** 10.64898/2026.06.30.735624

**Authors:** Adolfo A. Delgado, Alejandra Samano, Mahul Chakraborty

## Abstract

The maintenance of functional repeat arrays on nonrecombining sex chromosomes presents an evolutionary paradox: tandem repeats are intrinsically unstable yet must preserve sequence identity and copy number to remain functional. The Y-linked *Suppressor of Stellate* (*Su(Ste)*) locus in *Drosophila melanogaster* is a large tandem array that produces piRNAs to silence the X-linked meiotic driver *Stellate*, but how such arrays are maintained remains unclear. Here, we reconstruct and compare repeat-resolved assemblies of the *Su(Ste)/PCKR* tandem array across three strains and show that the array is partitioned into discrete domains of elevated sequence identity. These domains exhibit an alternating pattern of similarity, in which nonadjacent regions are more similar to each other than to neighboring regions, and this organization is conserved across strains. Copy-number variation occurs primarily within specific domains, while overall array architecture remains stable. These results indicate that concerted evolution in the *Su(Ste)* array operates within structurally defined domains rather than uniformly across the array. The association of domain boundaries with inverted repeat elements suggests that higher-order structure constrains gene conversion, shaping both sequence homogenization and copy-number dynamics. In contrast, Y-linked rDNA arrays show uniform sequence similarity across long genomic distances, indicating a distinct mode of homogenization. Together, our findings demonstrate that gene conversion on nonrecombining chromosomes is structured by higher-order array architecture, providing a general framework for the maintenance of functional repeat arrays.

## Introduction

Sex-linked genetic elements often arise due to sexually antagonistic selection, which favors alleles that enhance fitness in one sex while being neutral or deleterious in the other (Rice 1992). One consequence of this process is the accumulation of male- and female-benefiting genetic elements on heteromorphic sex chromosomes such as the Y and W, respectively, which are transmitted exclusively through a single sex. Studies of this process often focus on the acquisition and retention of translocated or duplicated protein-coding genes (Chakraborty et al. 2023). However, sexually antagonistic selection can also act on multi-copy genetic elements, particularly when the gene copy number itself is under selection (Bachtrog et al. 2019; Ellison and Bachtrog 2019; Martí and Larracuente 2023)

A fundamental challenge for such elements is that heteromorphic sex chromosomes typically lack meiotic recombination, reducing the efficiency of selection, accelerating the accumulation of deleterious mutations, and limiting the spread of adaptive alleles (Charlesworth and Charlesworth 2000; Bachtrog 2013). Tandemly repeated sequences may be especially vulnerable in this context due to their high intrinsic mutation rates (Flynn et al. 2018; Porubsky et al. 2025). However, intra-chromosomal gene conversion between duplicated elements provides a recombination-like mechanism on otherwise non-recombining chromosomes. Gene conversion can increase sequence similarity among repeats beyond what would be expected if they were evolving independently, a process known as concerted evolution. Through this process, duplicated sequences evolve in a coordinated manner, allowing functionally important repetitive elements to maintain sequence integrity over time (Connallon and Clark 2010).

Empirical support for this model comes from classical systems such as ampliconic regions and palindromes of mammalian Y chromosomes, where gene conversion maintains testis-specific gene families (Rozen et al. 2003; Rhie et al. 2023). While similar palindromic structures have been observed in other organisms, it remains unclear how large functional tandem arrays are structurally organized and maintained in diverse repeat-rich loci beyond well-characterized mammalian systems (Betrán et al. 2012). In these contexts, it is particularly unclear how gene conversion and concerted evolution operate across large arrays—whether sequence homogenization occurs uniformly across repeats or is spatially constrained by higher-order genomic structure. Addressing this question has been limited by the difficulty of assembling and comparing highly repetitive regions across individuals (Bachtrog 2013).

The Y chromosome of *Drosophila melanogaster* is largely heterochromatic, gene-poor, and nonrecombining (Kennison 1981; Bonaccorsi and Lohe 1991; Carvalho et al. 2001; Chang and Larracuente 2019). Despite this, it plays critical roles in male fertility, genome-wide regulation, and adaptation (Lemos et al. 2008; Larracuente and Clark 2013; Brown et al. 2020). Much of this functional impact derives from large blocks of repetitive DNA, including tandem arrays and rDNA clusters whose copy number and organization can vary substantially (Krsticevic et al. 2015; Brown et al. 2020; Martí and Larracuente 2023). These features make the Y chromosome a tractable model for studying the evolution and maintenance of functional repeat arrays on nonrecombining chromosomes.

A well-studied example of a Y-linked repeat array is *Suppressor of Stellate* (*Su(Ste)*), which suppresses an X-linked meiotic driver *Stellate* through piRNA–mediated silencing (Adashev et al. 2020). *Su(Ste)* originated from duplication of the autosomal gene *Ssl* and is closely related to both X-linked *Stellate* and the Y-linked paralog pseudo-*CK2*β repeats (*PCKR)* (Chang et al. 2022). The *Su(Ste)* tandem array evolved in response to the X-linked *Stellate*, which produces protein crystals that disrupt the development of Y-bearing sperm (Meng and Yamashita 2026). Because effective suppression requires sequence complementarity and sufficient piRNA production, both sequence identity and copy number of *Su(Ste)* are under selective pressure, creating a coevolutionary arms race (Palumbo et al. 1994; Aravin et al. 2001; Adashev et al. 2020).

Although the genetic interaction between *Stellate* and *Su(Ste)* has been extensively characterized, the physical architecture of the *Su(Ste)*/*PCKR* arrays and the mechanisms that maintain their integrity on a nonrecombining chromosome remain unclear (Chang and Larracuente 2019; Adashev et al. 2020; Chang et al. 2022). Here, we use improved, repeat-resolved assemblies of the *Su(Ste)*/*PCKR* array across multiple *D. melanogaster* strains to examine how concerted evolution and copy-number turnover shape a functional repeat array. We show that sequence homogenization is not uniform across the array but is instead partitioned into discrete domains defined by higher-order structure. We contrast this architecture with Y-linked rDNA arrays, which exhibit more globally distributed homogenization, revealing that concerted evolution can operate at distinct spatial scales even within the same chromosome.

## Results

### Assembly of Y-linked repeats in three strains

Characterizing variation in Y-linked repetitive regions such as *Su(Ste)* requires assemblies that accurately resolve their structural organization. However, the *Su(Ste)* locus remains fragmented in the existing Y assemblies, with different assemblies recovering complementary portions of the array (Chang and Larracuente 2019; Shukla et al. 2025; Carvalho et al. 2026) (Supplementary Fig. 1). Thus, combining these assemblies into a more contiguous framework is required to resolve the structure and evolution of Y-linked tandem arrays. To improve the resolution of the *Su(Ste)* region, we generated a more contiguous iso-1 assembly by merging existing assemblies using a repeat-aware approach (see Methods)(Chakraborty et al. 2016; Chang and Larracuente 2019). This assembly added ∼1.7 Mb of sequence relative to the iso-1 reference and substantially increased contiguity (contig N50 = 796,144 bp, scaffold N50 ∼2.5 Mbp), joining multiple previously fragmented regions, including loci near the *Su(Ste)* and rDNA arrays (Fig. 1A, B; Table 1). Uniform coverage of mapped HiFi reads across joins suggests that improved contiguity is not due to misjoins (Supplementary Fig. 1). Concordance with a previously published iso-1 Y assembly (Chang and Larracuente 2019) indicates consistency in large-scale structure (Fig. 1B). In addition, Nanopore direct RNA-seq data from an inbred strain A4 validates all 13 canonical Y-linked fertility genes in their expected locations (Supplementary Table 1), confirming gene-level accuracy. Notably, the locations of the *Su(Ste)* and rDNA arrays are consistent with prior cytological and deficiency maps, further supporting accurate recovery of Y chromosome organization (Carvalho et al. 2001; Hoskins et al. 2015).

**Figure 1.**
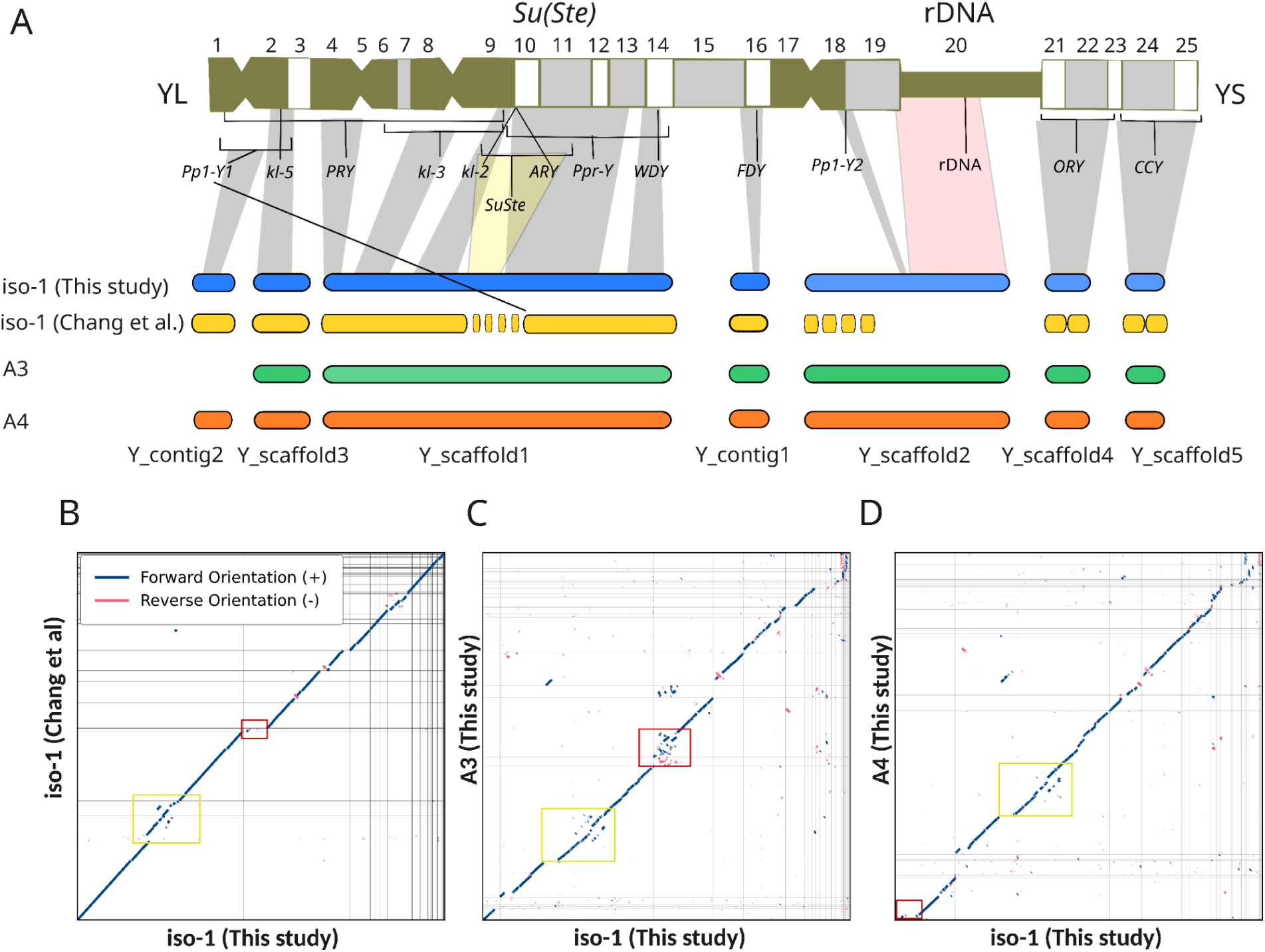
Y chromosome assembly of three *D. melanogaster* strains. **(A)** Schematic of the canonical cytogenetic map of the Y chromosome (h1–h25), with inferred positions of major genes and repeat arrays (Carvalho et al. 2000, 2001; Vibranovski et al. 2008; Hoskins et al. 2015). Below are shown assembled scaffolds and contigs for iso-1, A3, and A4 containing the fertility genes, illustrating the recovery of Y-linked regions, including the *Su(Ste)* (yellow) and rDNA (red) arrays. A previously published Y chromosome assembly (Chang and Larracuente 2019) is included to highlight improvements in contiguity relative to our iso-1 assembly **(B)** Dot plot alignments between the iso-1 Y assembly generated in this study and a previously published Y chromosome assembly (Chang and Larracuente 2019), showing concordance between the two assemblies. Dot plots between (**C**) iso-1 and A3 and (**D**) iso-1 and A4 Y assemblies, highlight conservation of gene order, orientation, and major Y-linked features across the strains.

**Table 1:** Contiguity statistics for Y chromosome assemblies.

| Assembly | # of scaffolds | Total size (bp) | Scaffold N50 | # of contigs | Contig N50 |
| --- | --- | --- | --- | --- | --- |
| **Y chromosome** |  |  |  |  |  |
| This study (iso-1) | 41 | 16,294,288 | 2,582,040 | 107 | 796,144 |
| Chang et al. (iso-1) | 55 | 14,603,684 | 686,238 | 80 | 416,887 |
| Release 6 (iso-1) | 13 | 4,162,520 | 3,667,352 | 261 | 81,922 |
| This study (A4) | 51 | 16,396,651 | 1,610,879 | 85 | 675,447 |
| This study (A3) | 37 | 17,034,472 | 2,762,414 | 57 | 831,042 |

To examine variation in Y-linked repeat arrays across strains, we scaffolded Y-linked contigs from the assemblies of the inbred strains A3 and A4 (Shukla et al. 2025) using the iso-1 assembly as a reference (Figs. 1C; D). These assemblies exhibit comparable contiguity to iso-1 (contig N50: A3 = 831,042 bp and A4 = 675,447 bp; scaffold N50: A3 = ∼2.76 Mbp and A4 = ∼1.61 Mbp; Table 1). A comparison of A3 and A4 assemblies against our iso-1 reference shows overall preservation of order and orientation of major Y-linked features across strains, including the major fertility genes and large ampliconic arrays, such as the *Su(Ste)* and rDNA arrays (Figs. 1C; D; Supplementary Figs. 2, 3).

### Structure of the *PCKR* and *Su(Ste)* locus

The *Stellate–Suppressor* of *Stellate* (*Su(Ste)*) system is a well-characterized example of sex-linked genetic conflict, in which an X-linked driver (*Stellate)* is suppressed by Y-linked piRNA-producing *Su(Ste)*. In the iso-1 assembly, *Su(Ste)* is organized as a single tandem array containing 358 repeat units, each of which contains a ∼1.1 kb *Hoppel/1360* element at the 5’ end and a 485 bp transcribed functional target domain (FTD) at the 3’ end (Fig. 2A, C). Within the array 203 *Su(Ste)* units carry a truncated *β*NAC*tes1* promoter sequence, which can potentially initiate transcription in the sense direction (Fig. 4A). Although *PCKR* has been proposed to localize proximal to *Su(Ste)*, its precise genomic position on the Y chromosome has remained unclear (Chang and Larracuente 2019). In our assembly, the *PCKR* repeats are located immediately adjacent to the 5’ end of the *Su(Ste)* array, between the fertility genes *ARY* and *kl-2* (Fig. 2A). Each *PCKR* unit carries a ∼1.3kb *HeT-A* non-LTR TE fragment at the 5’ end and a 251 bp sequence homologous to the FTD region of *Su(Ste)* at the 3’ end. Together, the *PCKR* and *Su(Ste)* arrays span ∼2.6 Mb, with *PCKR* occupying ∼0.19 Mbp and *Su(Ste)* ∼0.99 Mb, while the rest of the array consists of TE insertions (Fig. 2A, Supplementary Fig. 4). Although the two arrays are adjacent, the *PCKR* arrays across different genetic backgrounds show Hi-C contact mostly with the 3’ terminal of the *Su(Ste)* array, suggesting three-dimensional proximity between the two ends of the combined array (Fig. 2B, Supplementary Fig. 5).

**Figure 2.**
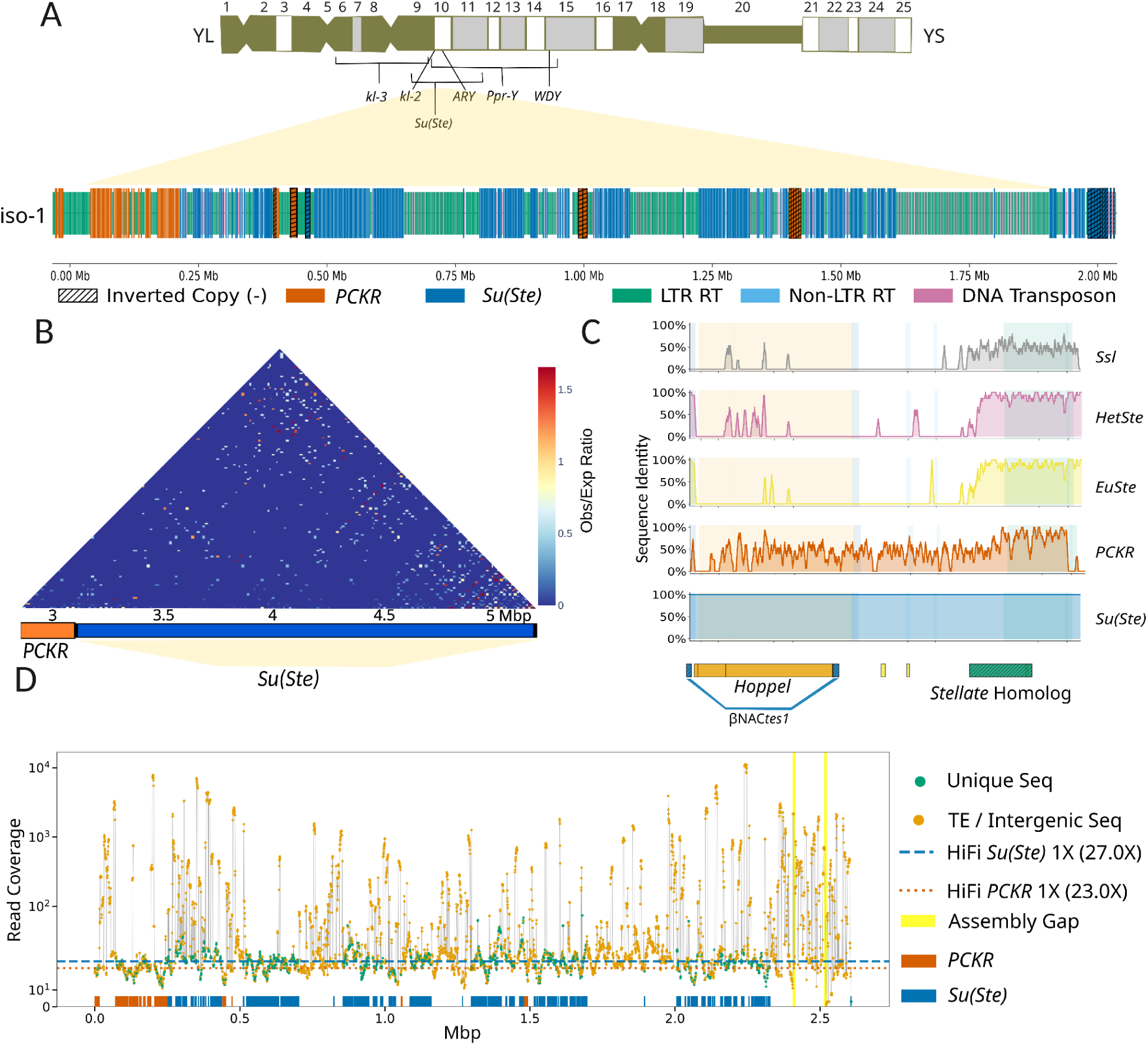
Organization of the *Su(Ste)* and *PCKR* arrays in iso-1. **(A)** Structural organization of the *PCKR*/*Su(Ste)* array in iso-1. **(B)** Hi-C contact map of Canton-S (Liu et al. 2025) sequencing reads mapped to the iso-1 *PCKR/Su(Ste)*, showing limited interactions between the two arrays. **(C)** Sequence similarity of *Ssl*, X-linked heterochromatic *Stellate* (*HetSte*), euchromatic *Stellate* (*EuSte*), and *PCKR* relative to the iso-1 *Su(Ste)* consensus. **(D)** HiFi sequencing read depth across the iso-1 *PCKR/Su(Ste)* array, supporting continuity and copy-number estimates. Dashed red and dotted blue lines denote the average coverage for non-repetitive sequences of *Su(Ste)* and *PCKR* units, respectively. Unique sequence refers to non-TE sequences within *PCKR/Su(Ste)* repeat units. The absence of assembly gaps within the main array supports the contiguity of our merged sequence.

Although the transcribed region of *Stellate* (functional target domain) shows homology with *PCKR*, *Ssl*, and *Su(Ste)* (Chang and Larracuente 2019; Adashev et al. 2020), we identified an uncharacterized gene *CG40635* located on an unplaced contig (211000022278760) in the iso-1 release 6 assembly (Hoskins et al. 2015), which exhibits sequence similarity with *Su(Ste)* and *PCKR* (Fig. 2C, Supplementary Figs. 6-8). The *CG40635* sequence is 98.4% identical to the terminal 251 bp of *PCKR* (position 2333-2584), suggesting that *CG40635* is likely a *PCKR* unit located in a fragmented contig (Supplementary Figs. 6-7). Interestingly, the 3’ region of *PCKR* (position 2113 to 2560) exhibits high sequence identity to the functional target domain (position 2123-2608) of *Su(Ste)*(97.4%), but less similarity to X-linked euchromatic (77.4%) and heterochromatic *Stellate* (78.5%) (Fig. 2C). This pattern indicates shared ancestry and partial functional conservation, with elevated similarity to *Su(Ste)* beyond that expected from shared ancestry alone (Fig. 2C). Notably, the *Su(Ste)* target region shows comparable similarity to both euchromatic (*Ste: CG33237*) and heterochromatic (*SteXh: CG42398*) paralogs (87.7% and 89.3%) respectively, suggesting that they are equally diverged from the X-linked *Stellate* sequences (Fig. 2C). In addition, a comparison of full *Su(Ste)* and *PCKR* consensus units shows that the *Hoppel* and *HeT-A* TE fragments within *Su(Ste)* and *PCKR* units, respectively, show 49.63% similarity (Fig. 2C). The dispersed sequence similarity observed between the TE fragments is likely due to shared short sequence motifs and not due to local exchanges as the reference *Hoppel* and *Het-A* shows similar sequence similarity (Supplementary Fig. 9).

### Read-Depth supports *PCKR* and *Su(Ste)* copy numbers

To determine whether copy number estimates for *PCKR* and *Su(Ste)* were affected by assembly errors, we mapped HiFi long reads from iso-1 to the unmasked array. To avoid inflated coverage from TEs shared across the genome, TE-derived sequences were excluded from coverage calculations, enabling read depth to be measured over the unique portions of repeat units. The non-repetitive sequences of *PCKR* and *Su(Ste)* exhibit similar continuous, uniform coverage (mean *PCKR*: 23×, mean *Su(Ste)*: 27×) (Fig. 2D). To ensure that these estimates were not biased by long-read sequencing, we performed the same analysis using paired-end Illumina data. Short-read coverage was similarly uniform as the HiFi coverage, although their mean coverage (*PCKR*: 59×, *Su(Ste)*: 64×) was nearly threefold greater than the HiFi coverage due to the higher genome-wide coverage of the Illumina sequence data (Supplementary Fig. 10). The concordance between HiFi and Illumina read coverage indicates that repeat copy numbers are accurately represented in the assembly, with no evidence of large-scale collapse or expansion (Supplementary Figs. 10 and 11). In addition, the average sequence divergence among the *PCKR* and *Su(Ste)* repeat units exceeds the error rates of HiFi reads, making misassembly unlikely despite their repetitive nature (Supplementary Table 2).

### Islands of concerted evolution in the *PCKR*/*Su(Ste)* array

To assess the extent of homogenization within the *PCKR* and *Su(Ste)* arrays, we performed pairwise sequence comparisons among repeat units in the iso-1 assembly. Pairwise sequence comparisons were performed using both gap-penalizing and non-penalizing alignment schemes. Because TE insertions can disproportionately influence similarity metrics in repeat-rich regions, we used the non-penalizing approach to better capture sequence similarity within the non-TE portions of *PCKR*/*Su(Ste)* repeat units, where localized gene conversion is expected to occur (Fig. 3A–D; Supplementary Fig. 12). Rather than uniform similarity, the *Su(Ste)* array is partitioned into eight discrete domains, hereafter referred to as “islands,” in which repeat units within each island are more similar to one another than to units in adjacent regions (Fig. 3A; Supplementary Figs. 13-15; Supplementary Table 3). Across the array, mean intra-island sequence identity is 98.00% (± 1.39%, n = 8,890), compared to 96.15% (± 0.97%, n = 13,096) between adjacent islands. This difference reflects a strong separation between intra- and inter-island similarity distributions (Mann–Whitney U test, *p* < 10×10^−300^) (Fig. 3E).

**Figure 3.**
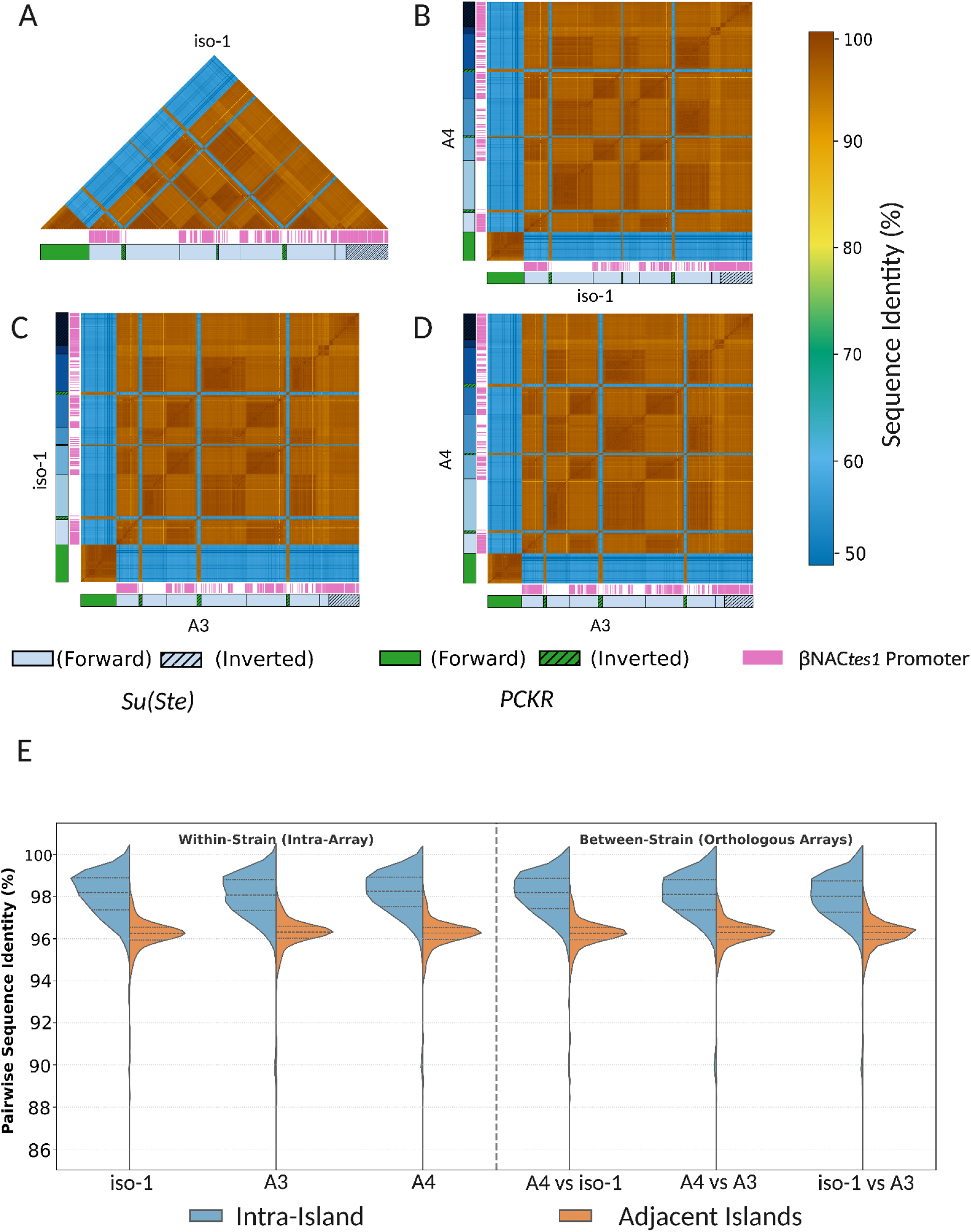
Higher-order architecture and sequence homogenization within the *PCKR/Su(Ste)* array. **(A)** Pairwise sequence similarity heatmap for iso-1, showing elevated identity within discrete domains (“islands”) and reduced similarity between adjacent islands. **(B-D)** Heatmaps comparing repeat units across iso-1, A3, and A4. Conserved patterns of higher-order architecture and sequence identity within and between corresponding islands demonstrate that array architecture is maintained across strains. **(E)** Sequence identity between *Su(Ste)* units within islands and between adjacent islands. Data is shown for both within-strain and between-strain comparisons.

These islands are arranged in an alternating pattern. Nonadjacent islands– particularly those with shared parity (odd- or even-numbered)–exhibit higher sequence similarity to each other than to neighboring islands. For example, Island 1 shares a greater identity with Islands 3 and 5 (∼97.4%) than with adjacent Island 2 (∼96.0%). Similarly, Island 3 is more similar to Island 5 (98.12%) than to Island 4 (96.48%), and comparable relationships are observed among even-numbered islands (Fig. 3A). Sequence identity drops sharply at island boundaries, which frequently coincide with inverted *PCKR* units (Supplementary Fig. 15), suggesting that these features may limit sequence exchange between adjacent domains. Notably, the terminal Island 8, composed entirely of inverted *Su(Ste)* units, shows reduced similarity to other islands, consistent with inverted units lowering homogenization rates (Nei and Rooney 2005). The islands also differ in sequence composition. The βNAC*tes1* promoter (Adashev et al. 2020) is unevenly distributed across the array, with enrichment in odd-numbered islands (70–96% of units) and reduced presence in even-numbered islands (0–41%) (Fig. 3A; Supplementary Table 4), suggesting differences in transcriptional potential across domains.

Although few inverted *PCKR* units occur within the *Su(Ste)* array, the *PCKR* array, in contrast to the multi-island structure of *Su(Ste)*, forms a single homogeneous domain, with high sequence identity across repeat units (∼98% in iso-1; Fig. 3A). Although *PCKR* units share substantial similarity with inverted *PCKR* copies embedded within the *Su(Ste)* array, overall sequence identity between the two arrays remains low (∼55%), consistent with limited inter-array homogenization. Together, these results reveal that the *Su(Ste)* array is not homogeneously structured but instead consists of discrete domains of concerted evolution, with homogenization occurring predominantly within islands rather than across the array as a whole.

### Islands of homogenization shared across strains

To determine whether the island structure and alternating sequence identity pattern are conserved across strains, we performed pairwise comparisons among repeat units across iso-1, A3, and A4 assemblies. Similar to iso-1, repeat units within the same island show higher sequence similarity than those in adjacent islands in both A3 and A4 (Fig. 3B, C, D; Supplementary Fig. 15). Mean intra-island identity across strains is ∼98.0%, compared to ∼96.2% between adjacent islands, and this difference is highly significant in all pairwise comparisons (p < 10^−300^, Mann-Whitney U test; Fig. 3E). This pattern is preserved not only within strains but also between them. Orthologous islands exhibit high sequence similarity across strains (e.g., ∼98% identity between corresponding islands in iso-1, A3, and A4). Moreover, the alternating sequence identity pattern is conserved across lineages: nonadjacent islands exhibit greater similarity than adjacent islands, even in inter-strain comparisons (Fig. 3B-D). For example, Island 5 in A4 shares a higher identity with Island 3 in iso-1 (98.12%) than with its adjacent island (∼96%), and similar relationships are observed among even-numbered islands. In contrast, comparisons between odd- and even-numbered islands consistently show reduced sequence identity (∼96%), reinforcing the distinction between alternating island groups. These patterns indicate that the relative positions and identities of islands have been maintained since the divergence of these strains. As concerted evolution is expected to increase divergence between haplotypes, the shared island structure between strains indicates a recent evolutionary origin. Together, these observations indicate that the eight-island organization and alternating homogenization pattern are evolutionarily young and stable features of the *Su(Ste)* array. The preservation of island boundaries—often marked by inverted *PCKR* units—and the consistency of sequence similarity patterns across strains suggest that these structural domains constrain sequence exchange over evolutionary timescales. This conservation suggests that higher-order array architecture imposes long-term constraints on sequence homogenization. Notably, the X-linked *Stellate* array shows no evidence of comparable domain-level partitioning (Supplementary Fig. 16), indicating that this structure is a specific property of the Y-linked *Su(Ste)* locus.

### Copy Number Variation within the *PCKR/Su(Ste)* Arrays

Because piRNA production from the *Su(Ste)* array is dosage-sensitive and may evolve in response to the copy number of the X-linked *Stellate* gene (Adashev et al. 2020), we examined variation in repeat copy number across strains. The *PCKR/Su(Ste)* arrays span 2.61, 2.79, and 2.98 Mb in iso-1, A3, and A4, respectively (Fig. 4A), indicating substantial variation in array size. Despite this variation, *PCKR* copy number is remarkably stable across strains (72–74 units), and unit structure is largely conserved, with most copies falling within a narrow size range (∼2.3–2.7 kb; Fig. 4A; Supplementary Table 5). In contrast, *Su(Ste)* copy number varies substantially, with iso-1, A3, and A4 carrying 358, 370, and 443 copies, respectively (Fig. 4A, B). Most *Su(Ste)* units fall within a relatively narrow size range (∼2.4–3.0 kb), with variation largely driven by insertions and deletions within associated TEs (Supplementary Fig. 17; Supplementary Table 5). Therefore, *Su(Ste)* copy number is an important contributor to the number of piRNA-producing functional units within the array.

**Figure 4.**
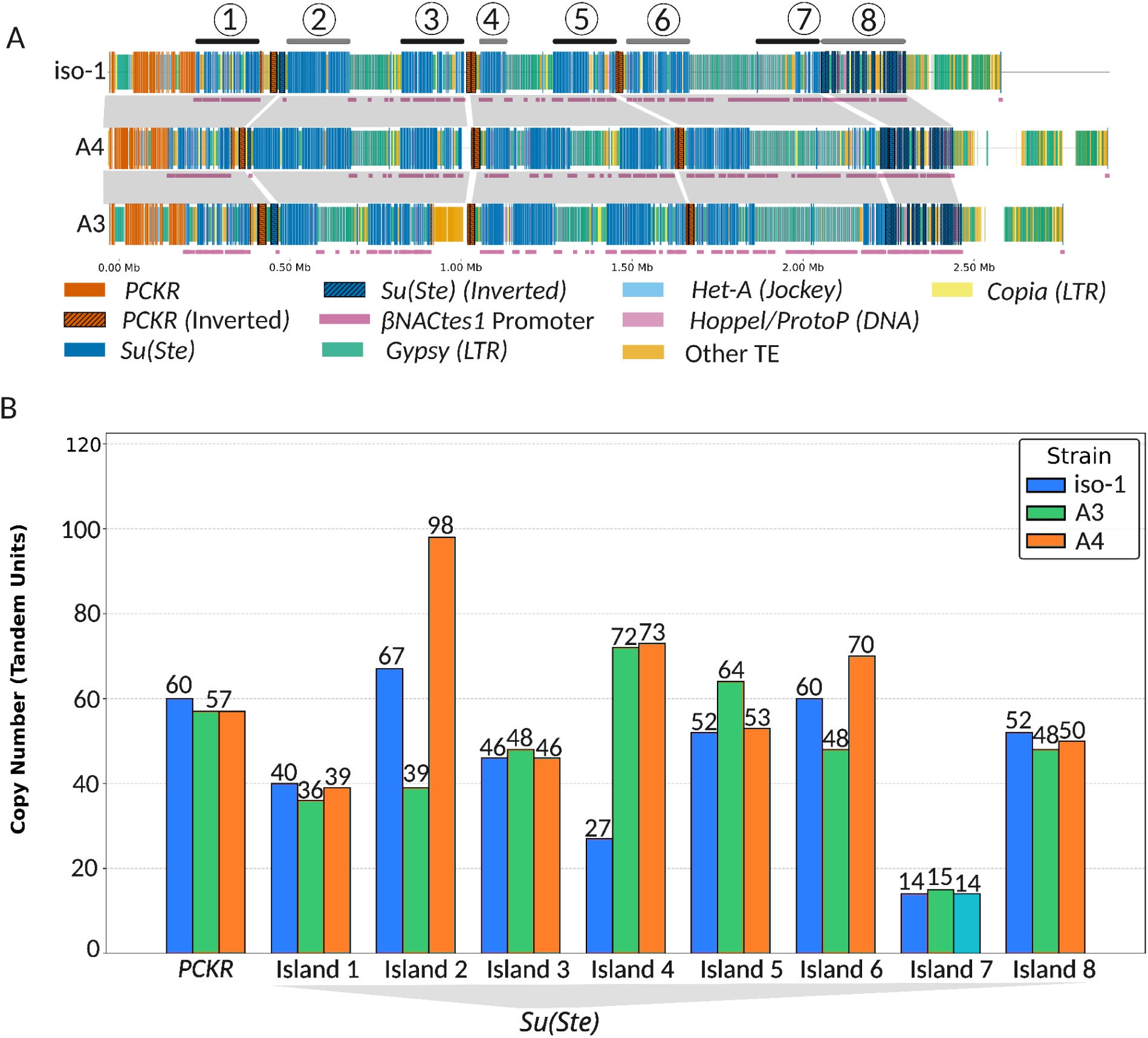
Copy Number Variation Within the Conserved Island Architecture of the *PCKR/Su(Ste)* array. (**A**) Comparison of syntenic regions of the *PCKR/Su(Ste)* arrays across iso-1, A4, and A3. Shaded regions indicate syntenic relationships between arrays inferred from conserved landmark anchors (inverted *PCKR* units). The size of the syntenic regions varies across strains. (**B**) Copy number of *PCKR* and *Su(Ste)* across islands in the three strains. Copy number variation of *Su(Ste)* is greater in specific domains within syntenic regions, particularly islands 2 and 6.

To determine how copy number variation is distributed across the array, we used conserved island boundaries—many of which coincide with inverted *PCKR* units—as landmarks (Shukla et al. 2025) to compare corresponding regions between strains. This analysis reveals that copy number variation is not uniform across the array but is concentrated within specific islands (Fig. 4 A, B). Terminal islands (e.g., Islands 1, 7, and 8) exhibit relatively stable copy numbers across strains (e.g., Island 1: 40, 36, and 39 units in iso-1, A3, and A4, respectively), whereas internal islands show substantial expansion or contraction (Fig. 4B). For example, Island 4 contains 27 copies in iso-1 but expands to 72 and 73 copies in A3 and A4, respectively, and Island 2 varies from 39 copies in A3 to 98 copies in A4. In contrast, other internal islands (e.g., Island 3) show relatively little variation across strains (46–48 copies) (Fig. 4B). Notably, islands that undergo the greatest expansions (e.g., Islands 2 and 4) are typically depleted of the βNAC*tes1* promoter, whereas more stable islands retain a high proportion of promoter-containing units (Fig. 4A). This pattern suggests that copy number dynamics vary systematically across the array and may be linked to local functional constraints.

These Y-linked copy number changes coincide with variation in X-linked *Stellate* copy numbers across the same strains. X-linked euchromatic *Stellate* arrays are massively expanded in A4 relative to iso-1 and A3 (iso-1: 11; A3: 2; A4: 198), whereas heterochromatic *Stellate* arrays are reduced in A4 (iso-1: 21; A3: 20; A4: 2;) (Supplementary Fig. 16; Supplementary Table 7)(Shukla et al. 2025). Together, these results suggest that copy number variation in the *Su(Ste)* array may evolve in the context of divergent *Stellate* dosage across strains.

### Contrasting patterns of concerted evolution in Y-linked rDNA units

To compare patterns of concerted evolution in *Su(Ste)* with another Y-linked tandem array, we examined sequence variation in the Y-linked rDNA array. Unlike *Su(Ste)*, the rDNA array is only partially assembled, with ∼1.12 Mb of rDNA sequence recovered at a scaffold terminus in iso-1 (Fig. 1A, Fig. 5A; Supplementary Fig. 18). The recovered iso-1 assembly contains 69 rDNA units, each of which contains the conserved rRNA coding sequences of 18S (1,995 bp), 5.8S (123 bp), 2S (30 bp), and 28S (∼3.9 kb) and variable non-coding sequences, including Intergenic Spacer (IGS), Non-transcribed Spacer (NTS) repeats (Supplementary Fig. 19). However, 28 of these also possess *R2* TE insertions, whereas 17 carry *R1* TE insertions, and one contains an I_dm TE (Fig. 5B, C) (Averbeck and Eickbush 2005). The units lacking TEs range from ∼11 kb to ∼20 kb, whereas units with TE insertions can reach up to ∼26.8 kb. A k-mer–based, assembly-free analysis (Rautiainen 2024) identified 18 major rDNA variants, 12 of which are present in the assembly, indicating that the assembly captures a representative subset of rDNA diversity (Supplementary Fig. 19).

**Figure 5.**
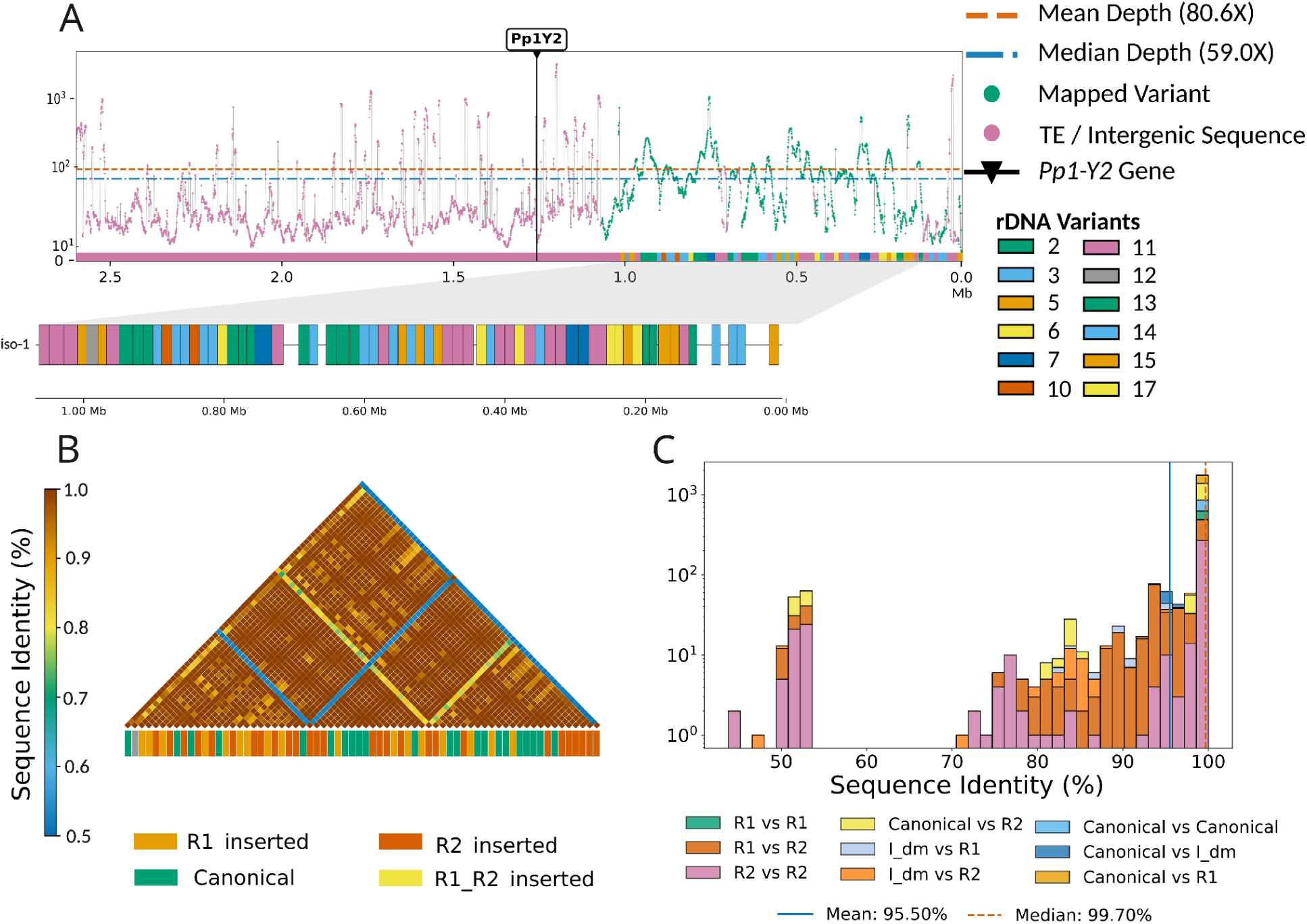
Homogenization patterns in Y-linked iso-1 rDNA arrays. **(A)** HiFi read-mapping coverage across the Y-linked rDNA-containing scaffold in iso-1. Ribotin-assembled variants mapped to the array highlight the variation present within the array (see Methods). **(B)** Heatmap of pairwise sequence identity between iso-1 Y-linked rDNA repeat units, categorized by transposon insertion status. **(C)** Distributions of sequence identity between different rDNA unit classes (Canonical, *R1*-inserted, *R2*-inserted). In contrast to the *Su(Ste)* array, rDNA units show uniformly high sequence similarity linearly across the array, lacking the higher-order architecture seen in *Su(Ste)*.

Despite incomplete assembly, rDNA units exhibit high sequence similarity. Pairwise identity between adjacent canonical (non-TE) units averages 99.92% (median 99.93%), and remains high even across units separated by more than 20 repeats (mean ∼99.73%, median ∼99.84%), indicating that sequence homogenization occurs over both short and long distances (Fig. 5B, C; Supplementary Fig. 20). High sequence similarity is also maintained among structurally heterogeneous units. rDNA copies containing *R1* or *R2* insertions show high identity relative to canonical units (*R1* vs. non-TE ∼99.7%; *R2* vs. non-TE ∼95.7%), as well as among themselves (*R1* vs. *R1* ∼99.7%; *R2* vs. *R2* ∼91.4%), indicating that TE insertions do not disrupt array-wide homogenization (Fig. 5C; Supplementary Table 8; Supplementary Fig. 21). Despite this overall homogeneity, three *R2*-inserted units carry secondary insertions of LTR and non-LTR elements, resulting in reduced sequence identity (55–75%) relative to the rest of the array (Fig. 5C; Supplementary Fig. 21). Similar patterns are observed in A3 and A4, where rDNA units retain high sequence identity across the array (mean ∼94–97%) (Supplementary Fig. 20). Together, these results indicate that rDNA arrays undergo global homogenization, whereas homogenization in *Su(Ste)* is spatially restricted to discrete domains. These differences suggest that the scale at which concerted evolution operates can vary substantially across tandem arrays, even within the same chromosome.

## Discussion

The maintenance of functional repeat arrays on nonrecombining sex chromosomes presents a fundamental challenge, as tandem repeats are inherently unstable yet must preserve sequence identity and copy number to maintain function (Bachtrog 2013; Martí and Larracuente 2023). The *Su(Ste)* array faces a particular challenge in this context, as it must maintain sequence identity and copy number so that piRNAs produced by *Su(Ste)* can effectively silence *Stellate*, even as *Stellate* copy number and sequence evolve on the X chromosome (Adashev et al. 2020). At the same time, the lack of recombination on the Y chromosome may promote the accumulation of deleterious mutations and slow the spread of beneficial variants, such as those that enhance piRNA production. Gene conversion, which can lead to concerted evolution or sequence homogenization among repeat units, provides a mechanism to purge deleterious mutations and spread beneficial mutations (Mano and Innan 2008; Connallon and Clark 2010; Chang and Larracuente 2019).

Our reconstruction of the Y-linked *Su(Ste)/PCKR* tandem array reveals that sequence homogenization is not uniform but instead occurs through a structured form of concerted evolution. The array is partitioned into discrete “islands” of elevated sequence identity that exhibit a conserved, alternating pattern across strains, indicating that this organization is maintained under selection. The similarity of intra- and inter-strain sequence identity suggests that this architecture is evolutionarily recent, as independent gene conversion would be expected to increase divergence between strains over time (Liao 1999). Thus, this organization may have evolved under positive selection and subsequently maintained across strains. Taken together, these observations suggest that the island architecture may help resolve the competing demands of stability and adaptability: by restricting gene conversion to localized domains, the array can maintain a sufficient number of highly similar repeat units required for effective piRNA production, while preserving sequence heterogeneity between domains that may facilitate adaptation to evolving *Ste* targets.

The island organization suggests that gene conversion on nonrecombining chromosomes can be spatially constrained. Classical models of repeat evolution, in which arrays are assumed to evolve through concerted or birth-and-death processes, treat repeat units as either forming a well-mixed population that homogenizes across the array or diverging independently (Nei and Rooney 2005). In contrast, the *Su(Ste)* array behaves as a set of localized conversion neighborhoods, within which sequence exchange is efficient but between which it is limited. The coincidence of island boundaries with inverted *PCKR* elements suggests that array architecture itself shapes these dynamics. Inverted repeats are known to promote double-strand breaks and homology-directed repair, raising the possibility that *PCKR* elements contribute both to the expansion of the array and to the delimitation of sequence exchange (Ramakrishnan et al. 2018; Watanabe et al. 2018). The alternating similarity among nonadjacent islands further indicates that both array history and chromatin topology may shape these dynamics. This pattern could arise from duplication of multi-copy blocks that preserve shared ancestry among subsets of islands, or from preferential gene conversion between regions brought into proximity in three-dimensional space, as suggested by elevated Hi-C interactions between nonadjacent loci (Willard and Waye 1987; Hallast et al. 2013; Haws et al. 2022). Together, these observations indicate that the effective units of concerted evolution are not only individual repeats, but also higher-order domains defined by array structure.

The modular organization of the *Su(Ste)* array parallels higher-order repeat (HOR) systems in centromeric satellites, in which multi-copy repeat units are homogenized into blocks, and structural boundaries limit the spread of sequence identity (Logsdon and Eichler 2022; Piri et al. 2026). Similar constraints have also been observed in ampliconic gene families on mammalian Y chromosomes, where gene conversion via palindromic sequences maintains sequence identity within defined architectural units (Rozen et al. 2003; Betrán et al. 2012; Hallast et al. 2013). The *Su(Ste)* locus represents a functionally distinct yet conceptually equivalent system, demonstrating that modular organization, coupled with constrained gene conversion, can maintain repetitive DNA in the absence of recombination. Within this framework, selection appears to act not only on total copy number, but also on the internal composition of the array: islands that are depleted of *β*NAC*tes1* promoters exhibit the greatest copy-number expansion, suggesting that functional requirements, such as the balance between sense transcription and antisense piRNA production, may shape which regions of the array are free to diversify.

In contrast, the Y-linked rDNA arrays exhibit a different mode of concerted evolution (Schlötterer and Tautz 1994; Averbeck and Eickbush 2005). Despite being only partially resolved, the recovered rDNA units show extremely high sequence identity across both short and long distances, with little evidence of discrete modular domains. This pattern is consistent with more globally distributed homogenization, likely driven by repair-associated recombination such as *R2*-induced double-strand break repair (Nelson et al. 2023), and strong purifying selection on coding sequences required for ribosome function. In contrast, *Su(Ste)* evolves under sexually antagonistic selection to produce piRNAs that silence *Ste*, a process that tolerates greater sequence variation (Aravin et al. 2001). Accordingly, sequence identity among the *Su(Ste)* units can be more relaxed than that of rDNA units, highlighting how differences in functional constraint and array architecture produce distinct homogenization regimes even within the same chromosome.

Taken together, these results show that the maintenance of functional repeat arrays on nonrecombining sex chromosomes depends not only on the presence of gene conversion but also on how that process is partitioned by higher-order structure. In the *Su(Ste)* locus, array architecture constrains both sequence exchange and copy-number dynamics, allowing the array to remain functionally stable while retaining evolutionary flexibility. These findings extend classical models of repeat evolution by showing that gene conversion is structured by genomic architecture, providing a general framework for understanding how functional repetitive DNA is maintained.

## Materials and methods

### Y chromosome assembly

To improve the iso-1 Y assembly, we merged the Y assembly from Chang & Larracuente (2019) as a query with Y-linked contigs from Shukla et al. as a reference, employing Quickmerge v0.3 (-hco 8.0 -c 3.0 -l 740929 -ml 5000)(Chakraborty et al. 2016). The assembly achieved a contig N50 of ∼2.5 Mb but lacked the *Pp1-Y1* gene and several exons from *kl-3* (exons 2–4, 14, 15), *PRY*, *kl-5*, *ORY*, and *Ppr-Y*, attributed to HiFi sequencing or assembly bias (Carvalho et al. 2026). We manually curated the missing sequences (details below), resulting in a more complete Y assembly, though this introduced new gaps and reduced the contig N50 to 796 kb. The A3 Y-linked contigs from Shukla et al. (2025) were scaffolded using RagTag v2.1.0 (Alonge et al., 2019) with our new iso-1 assembly as the reference. The A4 Y-linked contigs were scaffolded using the A3 assembly as the reference, as it yielded a more contiguous assembly than using iso-1.

### Assembly validation

To assess assembly accuracy across the Y assemblies and at the joins from quickmerge, we mapped HiFi raw reads to the merged assembly using Winnowmap2 (v2.03) with repeat-aware parameters (k=28 and default PacBio parameters) (Jain et al. 2022) and a k-mer library from Meryl v1.4.1 (Rhie et al. 2020). Coverage was calculated with SAMtools v1.21 (samtools depth -a) in 2 kb windows, identifying regions with zero primary read coverage via BEDTools v2.31.1(Quinlan and Hall 2010), filtering for primary mappings. Regions with zero read coverage represent assembly gaps from the query assembly by Chang et al. (2019) or sequences lacking HiFi read coverage. The alignment file was further inspected in Integrative Genomics Viewer (IGV) (Thorvaldsdóttir et al. 2013) to confirm that long reads span the merging breakpoints.

To confirm the location and presence of the known major Y-linked features, including the fertility genes, the new iso-1 assembly was aligned to the previous assembly (Chang and Larracuente 2019) using Winnowmap2 (-x asm20). The A3 and A4 Y chromosome assemblies were aligned to our new iso-1 assembly as well. The resulting PAF alignment files were uploaded to the D-GENIES server (Cabanettes and Klopp 2018) to visualize the order and orientation of the contigs using a dot plot and to examine alignment across the major fertility genes and ampliconic arrays. The D-GENIES PAF files were also visualized using a custom python script utilizing plotly. Coverage analysis was performed using the same methods. The N50 and assembly size calculations, as well as gap identifications, were conducted using QUAST v5.3.0 (Gurevich et al. 2013).

### Manual Curation of Y-Linked Loci

To assess gene completeness, we mapped consensus coding sequences from Carvalho et al. (2026) onto each assembly using BLAT v3.7 (Kent 2002) and annotated them with a custom Python script (CDSreconstruction.py). We identified missing genes and exons and employed three strategies to recover them. For the *Pp1-Y1* locus, raw HiFi reads were mapped to the reference *Pp1-Y1* sequence and assembled with Flye v2.9.5 (Kolmogorov et al. 2019). The resulting contig contained highly repetitive termini, preventing scaffold integration, and was appended to the final assembly as Y_Contig2. For exon-level dropout within assembled scaffolds, as observed in *kl-3* (exons 2–4, 14, 15) and the *PRY* coding sequence, we conducted manual patching guided by synteny. Missing exon sequences and their flanking regions were extracted from the Chang et al. (2019) assembly and inserted into syntenic positions within the iso-1 Y scaffold. For additional missing exons in deep satellite arrays, particularly in *kl-5*, *ORY*, and *Ppr-Y*, we incorporated fully assembled contigs from Carvalho et al. (2025).

### Direct RNA sequencing and transcriptomic validation

To verify the structures and transcripts of Y-linked genes, including manually curated genes, we performed Oxford Nanopore direct RNA sequencing (dRNA-seq) on whole bodies of A4 male and female *D. melanogaster*, following our established protocol (Samano et al. 2026). Total RNA was extracted from 7-day-old flies using TRIzol and reverse transcribed with the Induro Reverse Transcriptase kit. Libraries were prepared using the Oxford Nanopore Direct RNA Sequencing kit (SQK-RNA004) and sequenced on a PromethION RNA flow cell for 72 hours. We also used published dRNA-seq data from the strain BL156 (Samano et al. 2026). The dRNA-seq data from A4 and BL156 flies were mapped to our ISO1 assembly using minimap2 (-ax splice) (Li 2018), and exons were classified as supported if they had an average mapping depth of at least one read (Supplementary Table 1). To differentiate Y-linked male-specific expression from autosomal cross-mapping artifacts (e.g., at the *FDY*/*vig2* paralogous loci), we mapped A4 female dRNA-seq data to the ISO1 Y assembly under the same parameters. Finally, we verified the coding sequences and exon-intron boundaries by visually inspecting read alignments and splice junctions at the targeted insertion boundaries using IGV.

### Repeat Annotation

Repetitive sequences were annotated using RepeatMasker v4.1.8 (Smit et al. 2013) with a custom repeat library (Khost et al. 2017). To improve annotation specificity, the library was divided into TEs and satellite sequences, and each assembly was annotated separately in two passes (TE-only and satellite-only). Simple repeat annotations produced by the integrated TandemRepeatFinder (TRF) module were further refined using a custom Python script (ComparativeRepeats.py). This script reconciled fragmented and reverse-complemented TRF annotations (e.g., AGAGA and CTCTT) by grouping them under their canonical cytological repeat name (e.g., AAGAG), thereby ensuring accurate abundance quantification.

### Structural annotation of *PCKR/Su(Ste)* array

To characterize the *Su(Ste)* and *PCKR* tandem repeat arrays, consensus sequences for both repeat families were mapped against the target genome assemblies using minimap2 (v2.29). Repetitive k-mer filtering was disabled to capture all copies within the array, retaining all mapping chains with parameters (-f 0, -P). Only alignments with a minimum target length of 1,850 bp and a sequence identity of 80% were considered, with the identity calculated relative to the mapped portion of the query sequence to account for large TE insertions. Following locus splitting, we computed copy numbers, average sequence identities, and strand orientations for each family. We investigated the homology targets of *PCKR*, *Su(Ste)*, *Stellate* (*EuSte* and *HetSte*), and *Ssl* loci in the *D. melanogaster* genome using the blastn algorithm from the BLAST+ toolkit (v2.16.0), utilizing the latest release of the *D. melanogaster* reference genome gene regions and transcripts (release 6.67)(Hoskins et al. 2015). Alignments were limited to the top five transcript hits for each query sequence. The βNAC*tes1* sequence within the array was annotated by mapping it to the array using blastn with an E-value threshold of 1e-10.

To identify syntenic inverted *PCKR* units nested within the *Su(Ste)* arrays of the iso-1, A3, and A4 strains, we mapped the *PCKR* units to the *Su(Ste)* arrays using minimap2, retaining only negative-strand hits. Each inverted unit’s spatial position was assigned a relative coordinate (0.0 to 1.0) based on its center point relative to the overall span of the array. Inverted elements were deemed orthologous across the three strains if their relative positions differed by no more than within ±5%. Orthologous inverted *PCKR* unit sequences were aligned using MUSCLE (v5.1.0) to calculate sequence divergence.

### Sex-specific transcription of *PCKR/ Su(Ste)*

To characterize the male-specific transcripts from the *PCKR* and *Su(Ste)* tandem repeats, A4 male and female dRNA-seq reads were mapped to iso-1 *PCKR* and *Su(Ste)* consensus sequences using minimap2 (-ax splice -uf -k14) (v2.29). Resulting alignments were filtered and sorted using SAMtools (v1.21) using parameters -F 2308 -q 60 to exclude unmapped, secondary, and supplementary alignments. The male and female read coverage was calculated for each nucleotide using samtools depth.

### Comparative and phylogenetic analysis

To evaluate homology among array units, nucleotide sequences of the *PCKR* and *Su(Ste)* repeat units were aligned using MAFFT (v7.526) (Katoh and Standley 2013) with parameters --localpair --maxiterate 1000 to accommodate structural variations and TE insertions. Pairwise sequence identity was computed by dividing the total number of nucleotide matches by the alignment length, excluding positions containing gap characters in either sequence. A maximum-likelihood phylogenetic reconstruction of the iso-1 array units was conducted using IQ-TREE2 (v2.3.6)(Minh et al. 2020). The optimal nucleotide substitution model was identified with the integrated ModelFinder algorithm (-m MFP). To assess branch and node support, 1,000 ultrafast bootstrap (UFBoot) replicates (-B 1000) were performed, with nearest-neighbor interchange optimization to minimize the risk of overestimating branch supports due to sequence divergence or model violations (-bnni).

Insertion-deletion variation in the repeat units across the *PCKR* and *Su(Ste)* arrays was generated using a Python pipeline (InDel_Variation.py). A consensus sequence within the MAFFT multi-strain alignment was identified by masking columns with >50% gaps across all strains. For each strain, sequence variation profiles were calculated across all repeat units at every consensus nucleotide position, capturing: deletions (% of units missing sequence), divergence (% of units with SNPs/mismatches), and insertions (% of units containing unmasked characters).

### X-linked *Stellate* consensus

The X-linked *Euchromatic* (*EuSte*) and *Heterochromatic Stellate* (*HetSte*) units in the X chromosome were identified using blastn (BLAST+ v2.16.0; -dust no) with *EuSte* and *HetSte* consensus queries from Shukla et al (2025). Resulting alignments were filtered to retain hits with ≥90% sequence identity and >800 bp in length, ensuring that redundant alignments were excluded. The sequences of all *Stellate* units from all strains were pooled and aligned using MAFFT (v7.526) with the L-INS-i iterative refinement algorithm (--localpair --maxiterate 1000).

### Hi-C interactions

Adapter sequences from raw paired-end Hi-C sequencing reads originating from both the Canton-S (Liu et al. 2025) and iso-1 (Schauer et al., 2017) strains were trimmed, retaining reads with a minimum length of 35 bp. The Hi-C contact matrices were constructed using the Juicer pipeline (v1.6) (Durand et al. 2016) and iso-1 Y from this study as the reference, and were converted using the Cooler suite (Abdennur and Mirny 2020). Observed/Expected (O/E) ratios were calculated using the hicTransform tool of HiCExplorer (Ramírez et al. 2018). The contact matrix corresponding to the repetitive *Su(Ste)* and *PCKR* locus of the iso-1 Y chromosome (coordinates: 2,852,210–5,180,375 bp) was extracted, and the number of interactions between islands was quantified.

### *Su(Ste)* and *PCKR* copy number estimation

To evaluate the structural integrity of the *Su(Ste)* and *PCKR* arrays in the three strains, HiFi long reads were mapped to their respective unmasked array sequence using Winnowmap2, disabling secondary alignments (--secondary=no) and excluding supplementary alignments (-F 2308) to filter multi-mapping and prevent the double-counting of mapped reads. TEs in the array sequences were annotated with RepeatMasker (-nolow). Read depths per nucleotide were computed from the filtered BAM alignments using samtools depth (-a). To eliminate mapping artifacts caused by reads mapping inside TEs, read depths of sequences located within RepeatMasker-generated coordinates were excluded. We mapped Illumina paired-end short-read data from the same strains to their respective repeat-masked arrays using bwa-mem2 (Vasimuddin et al. 2019), excluding secondary and supplementary alignments (-F 2308) to reduce multimapping.

### Annotation of rDNA assembly

We used RepeatMasker annotations of the main rDNA-containing scaffold from each assembly to identify individual rDNA repeat units and their structural variants. The rDNA units were identified using the 18S rRNA sequence. For each 18S feature, the nearest 28S and ETS annotations establish unit boundaries and strandedness, ensuring each unit spans from the 18S start coordinate to the terminal ETS boundary. Units were then examined for the presence of non-LTR retrotransposons, including *R1*, *R2*, *Doc*, and *I_DM* elements. Units lacking internal insertions were classified as canonical, while disrupted units were tagged with their specific TE burden (e.g., *R1*-inserted or *R2*-inserted). To facilitate analysis of full units, we filtered out truncated or incomplete copies using a length filter, retaining units between 9,000 bp and 30,000 bp (Stage and Eickbush 2007).

### Assembly-free rDNA variant identification

We used Ribotin (Rautiainen 2024) to reconstruct the rDNA structural variants (morphs) of the iso-1 rDNA array. Specifically, we ran ribotin-ref using HiFi reads mapped to the iso-1 rDNA assembly and a single canonical rDNA unit. We mapped HiFi reads to the rDNA array using Winnowmap2 with the `map-pb` parameter and kept only primary alignments by filtering out secondary and supplementary alignments (samtools view -F 2308). Because accurate resolution of SVs requires spanning structurally variable spacer regions and internal TEs, we excluded reads shorter than 10,000 bp. We subsampled 30X of the total mapped reads (calculated using an array size of 1.05 Mb), which provided optimal recovery of rDNA variants. We used specific clustering parameters to differentiate highly similar spacer variants: approximate morph size was set to match the average unit size present in each assembly, with edit distance threshold of 10 for initial clustering (--morph-cluster-maxedit) and 1 for sub-clustering (--morph-recluster-minedit). To address complex local tangles in the assembly graphs, often induced by inverted TE sequences and negative-strand copies, we used a *k*-mer size of 101 for iso-1 during the MBG graph-building step. This size was adopted as the lowest functional threshold because larger k-mer sizes inflated error tolerances, leading to increased node sizes in the MBG graphs and the merging of distinct biological variants into artificial consensus sequences. To examine the proportion of Ribotin-assembled morphs present in the assembly, we mapped the morphs back to the iso-1 scaffolds using Minimap2 (v2.29). To ensure highly specific variant placement while accounting for minor sequencing or assembly errors in the contiguous scaffolds, alignments were processed with low (5% or -x asm5) and high (20% or -x asm20) sequence divergence parameters. Overlapping alignments were addressed by ranking alignment hits by Minimap2 alignment score (AS:i tag), ensuring that each unit in the assembly was annotated with its single best-matching rDNA variant.

### Homogenization patterns of rDNA arrays

We extracted assembled ribosomal DNA (rDNA) units from Y-linked arrays and classified sequences by insertion status (canonical versus transposable element-inserted) and by their position within the contiguous array. Alignments were performed using MAFFT (v7.526) with the FFT-NS-2 algorithm, optimized through iterative refinement parameters (--retree 2 --maxiterate 1000 --ep 0.0) to maximize alignment accuracy. To address the outsized impact of unique indels on pairwise divergence metrics, gap characters present in only a single repeat unit within the alignment matrix were heavily penalized. For each pairwise sequence comparison, sequence identity was calculated as the number of matching nucleotides divided by the total alignment length.

## Data availability

The iso-1 merged assembly and dRNA-seq data are available at NCBI Bioproject PRJNA1471322. All scripts and the scaffolded A3 and A4 Y assemblies are available at https://github.com/chakrabortymlab/Y3VAR.

## Acknowledgments

We thank Rich Meisel and Heath Blackmon for their comments on the manuscript. AD and MC were supported through the STEGG-INTERACT program funded by the National Science Foundation grant 2319694 to RA Zufall and RP Meisel. This work was also supported by the Texas A&M startup funds and the NIH grant R00GM129411 to M.C.

