## Supplemental Materials for "Higher-order Architecture Shapes Concerted Evolution in a Y-linked repeat array"

Affiliations:
Department of Biology, Texas A&M University, College Station, TX 77843

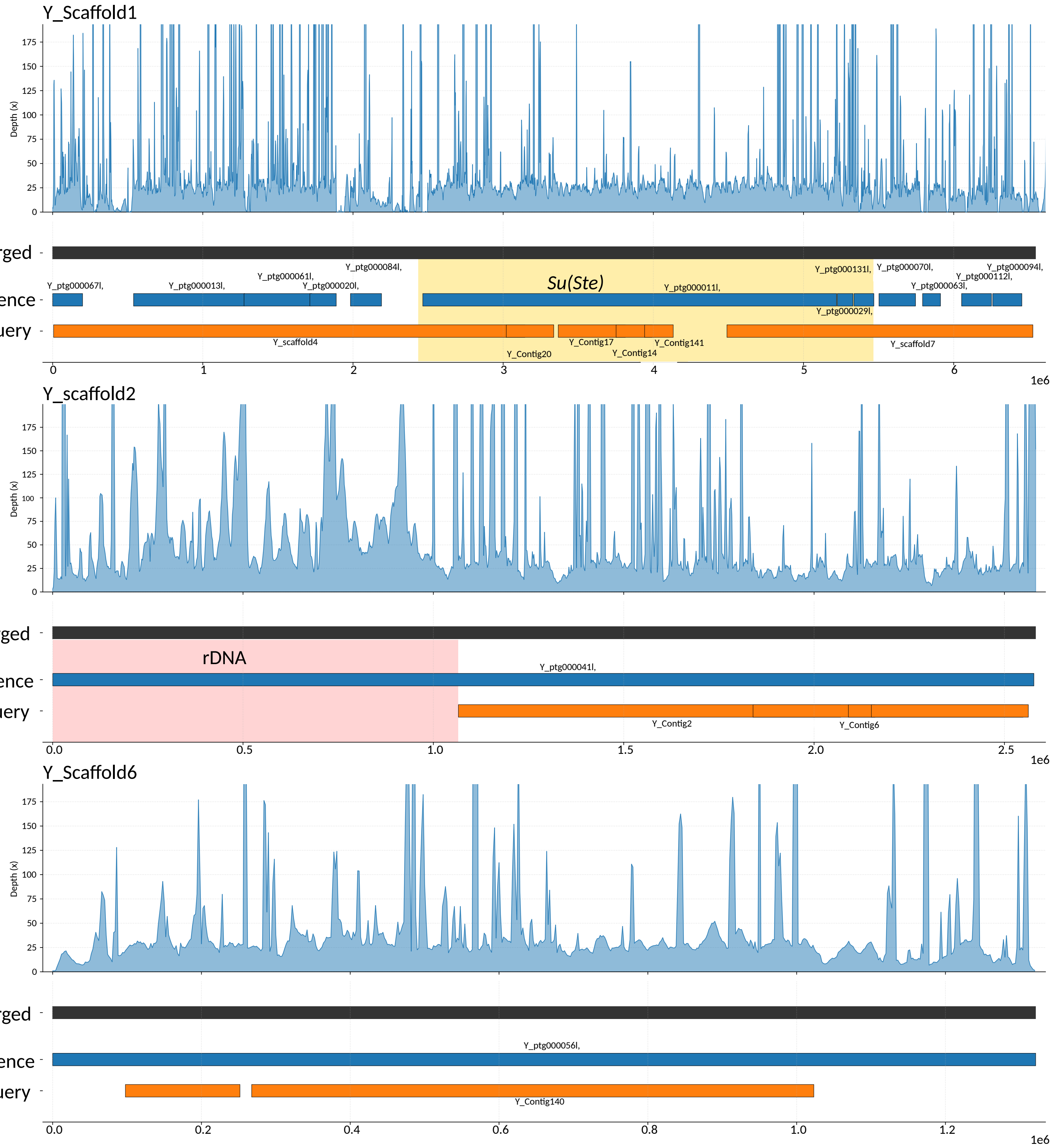

Y_Scaffold1
Depth (x)
Merged
Reference
Query
Su(Ste)
Y_ptg000067l,
Y_ptg000013l,
Y_ptg000061l,
Y_ptg000020l,
Y_ptg000084l,
Y_ptg000011l,
Y_ptg000131l,
Y_ptg000029l,
Y_ptg000070l,
Y_ptg000063l,
Y_ptg000112l,
Y_ptg000094l,
Y_scaffold4
Y_Contig20
Y_Contig17
Y_Contig14
Y_Contig141
Y_scaffold7
1e6
Y_scaffold2
Depth (x)
Merged
Reference
Query
rDNA
Y_ptg000041l,
Y_Contig2
Y_Contig6
1e6
Y_Scaffold6
Depth (x)
Merged
Reference
Query
Y_ptg000056l,
Y_Contig140
1e6

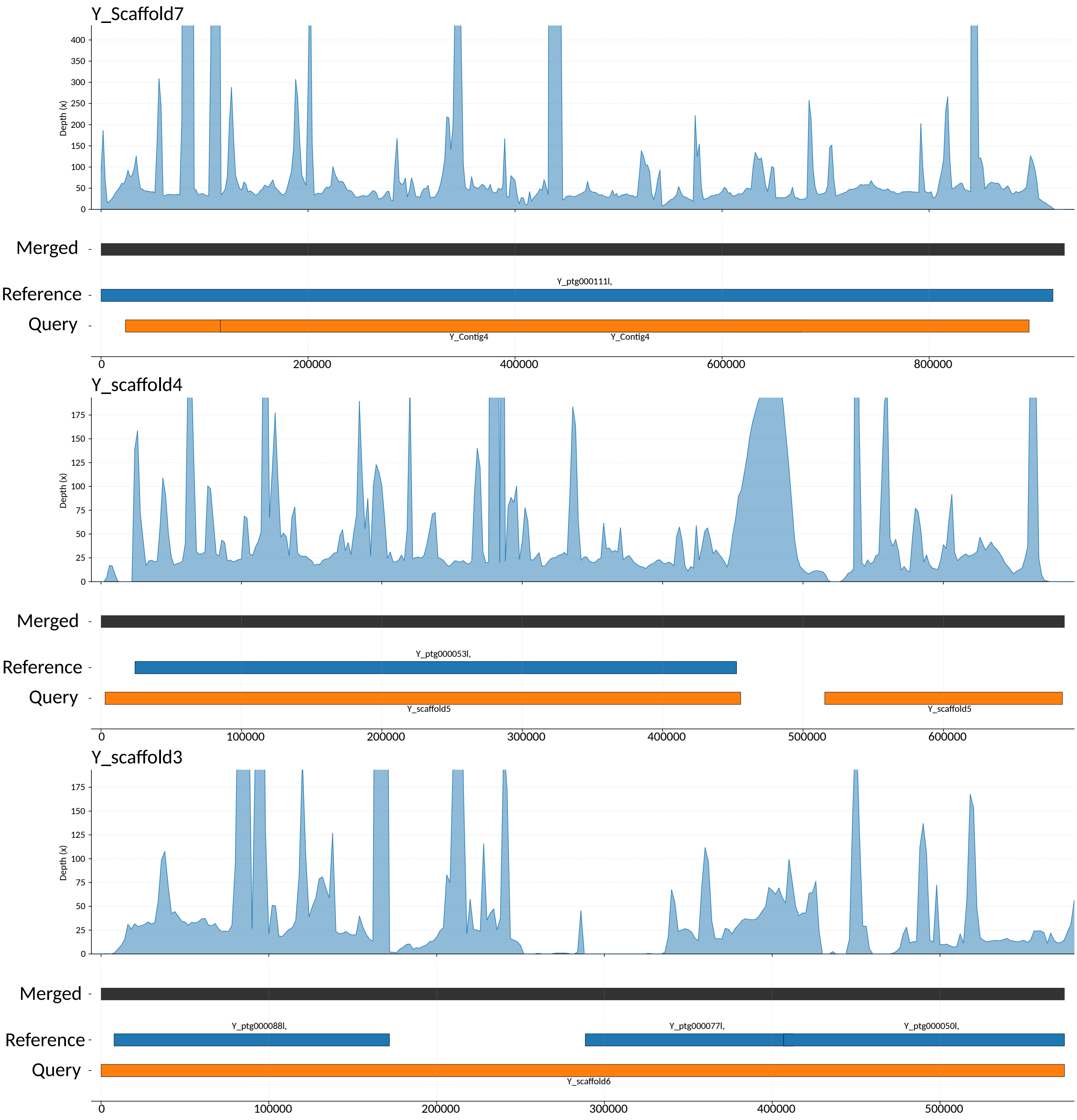
Y_Scaffold7
Depth (x)
0
50
100
150
200
250
300
350
400
Merged
Reference
Query
Y_ptg000111l,
Y_Contig4
Y_Contig4
0
200000
400000
600000
800000
Y_scaffold4
Depth (x)
0
25
50
75
100
125
150
175
Merged
Reference
Query
Y_ptg000053l,
Y_scaffold5
Y_scaffold5
0
100000
200000
300000
400000
500000
600000
Y_scaffold3
Depth (x)
0
25
50
75
100
125
150
175
Merged
Reference
Query
Y_ptg000088l,
Y_ptg000077l,
Y_ptg000050l,
Y_scaffold6
0
100000
200000
300000
400000
500000

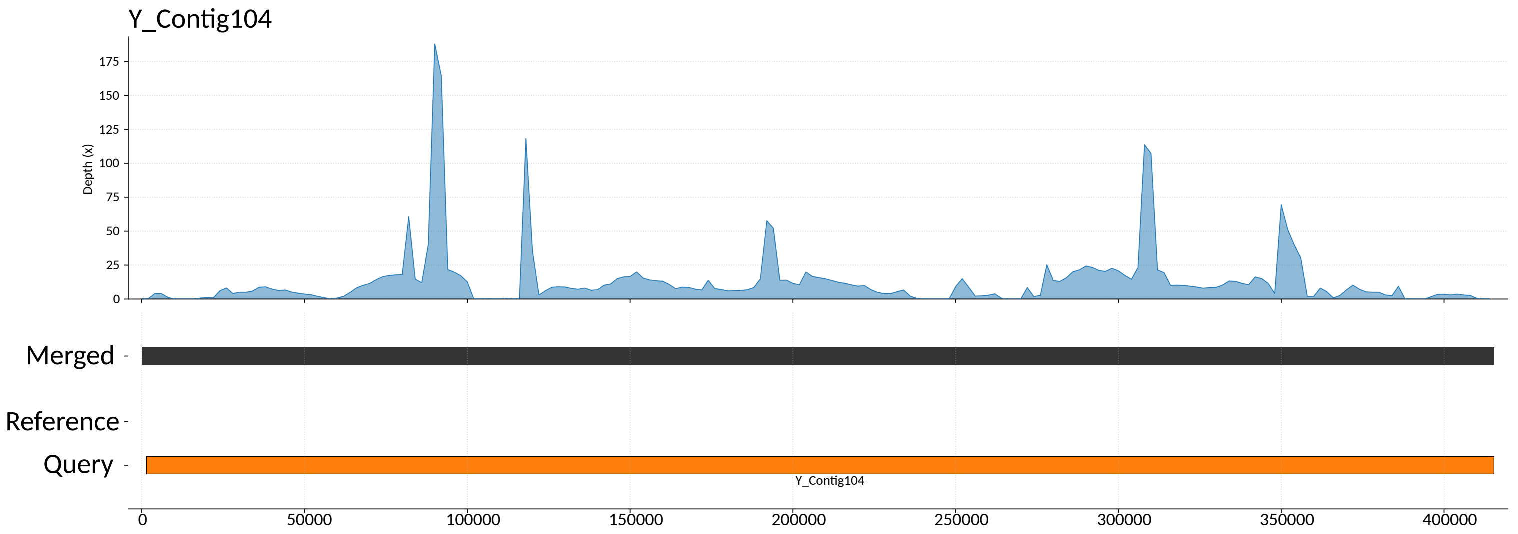
Y_Contig104
Depth (x)
0
25
50
75
100
125
150
175
Merged
Reference
Query
Y_Contig104
0
50000
100000
150000
200000
250000
300000
350000
400000

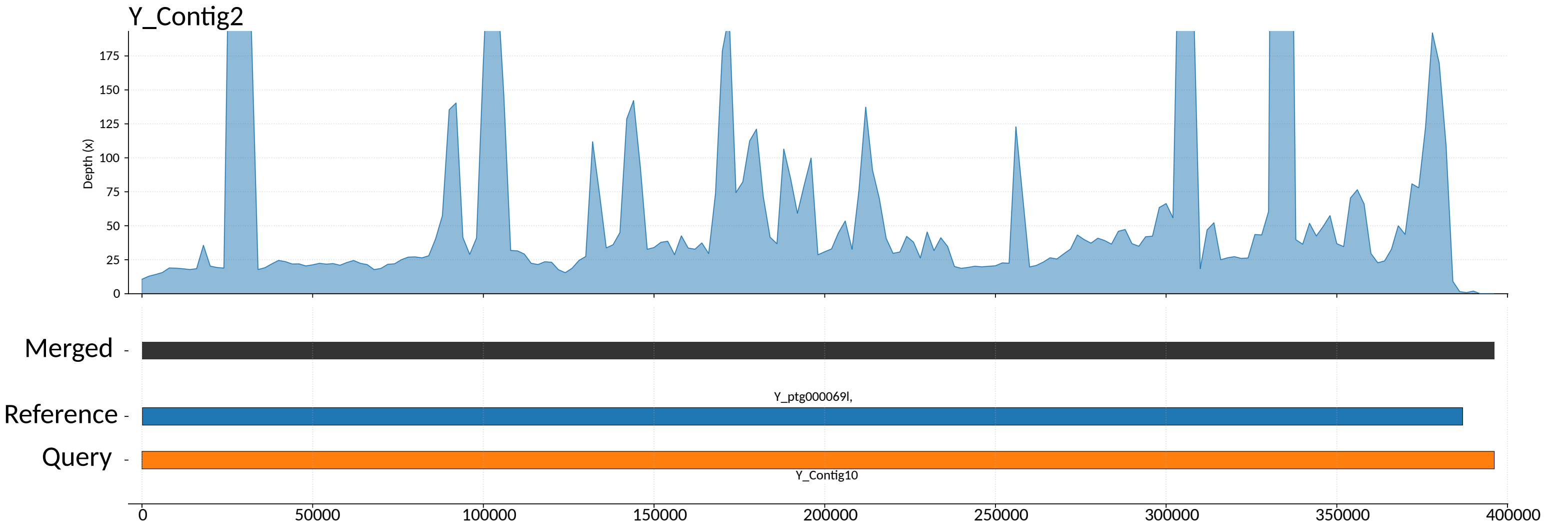
Y_Contig2
Depth (x)
0
25
50
75
100
125
150
175
Merged
Y_ptg000069l,
Reference
Query
Y_Contig10
0
50000
100000
150000
200000
250000
300000
350000
400000

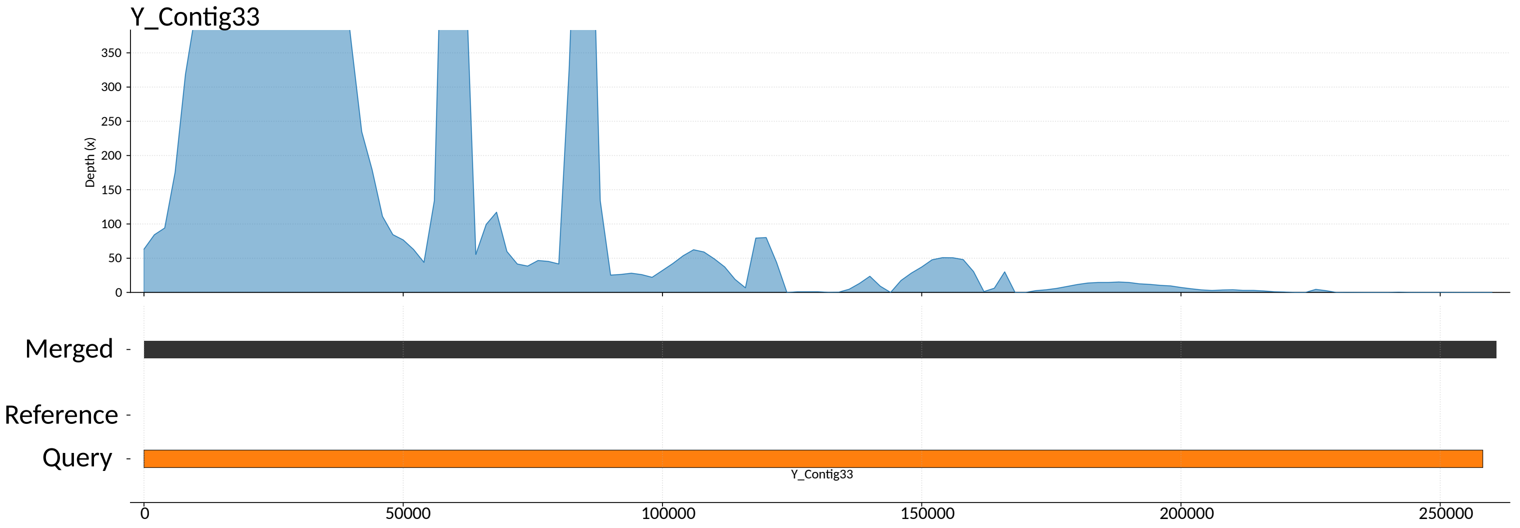
Y_Contig33
Depth (x)
0
50
100
150
200
250
300
350
Merged
Reference
Query
Y_Contig33
0
50000
100000
150000
200000
250000

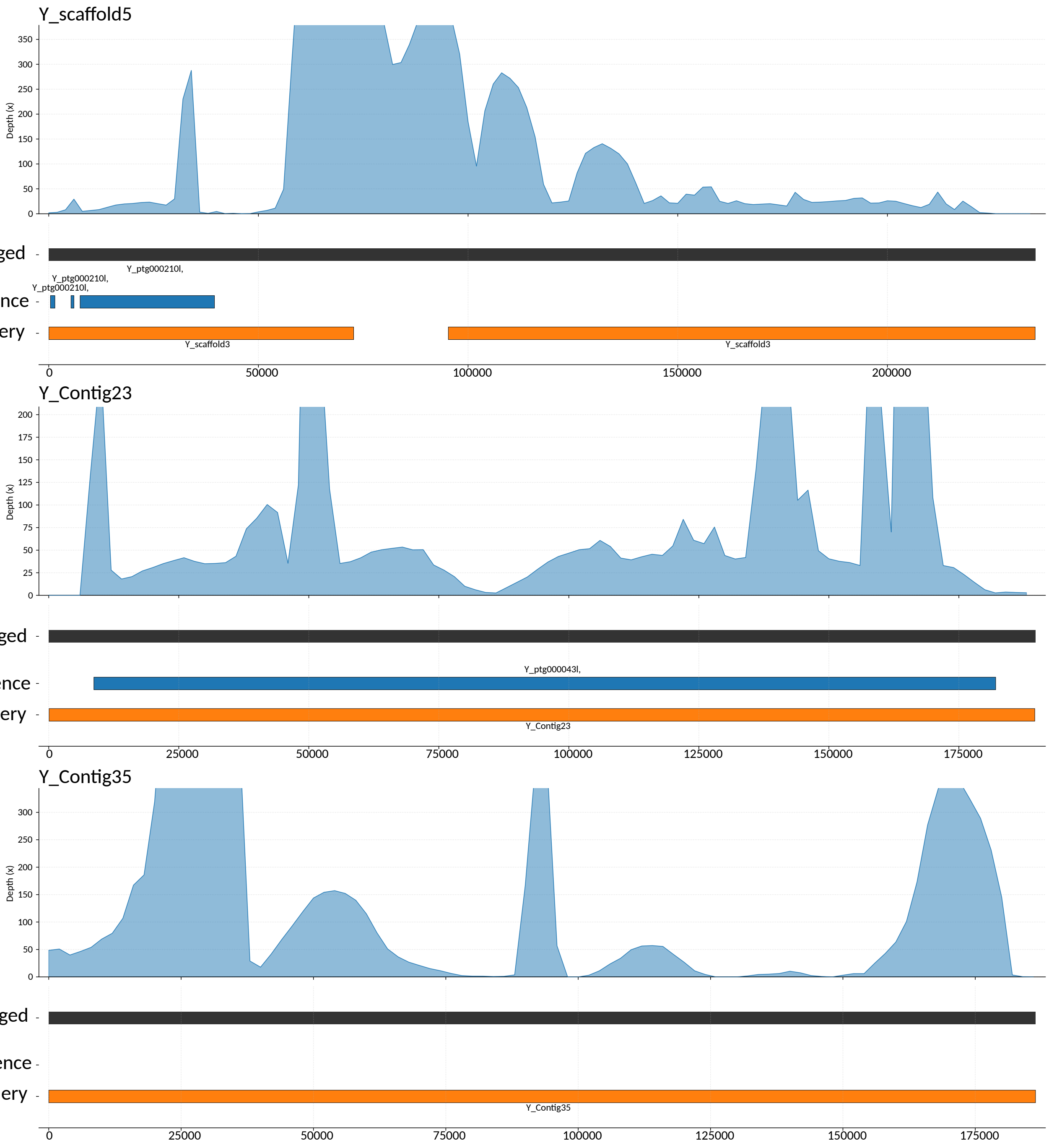

Supplementary Figure S1. Assembly merging and HiFi read coverage of the iso-1 Y chromosome. The final merged assembly (black) was constructed using the (Shukla et al. 2025) assembly (blue) as the reference and the (Chang and Larracuente 2019) assembly (orange) as the query. Regions from the reference are retained where supported, while the query sequence extends gaps based on overlapping anchors. Coverage profiles above each scaffold show read support across the merged assembly; localized drops in coverage correspond to regions lacking HiFi read support in the query assembly.

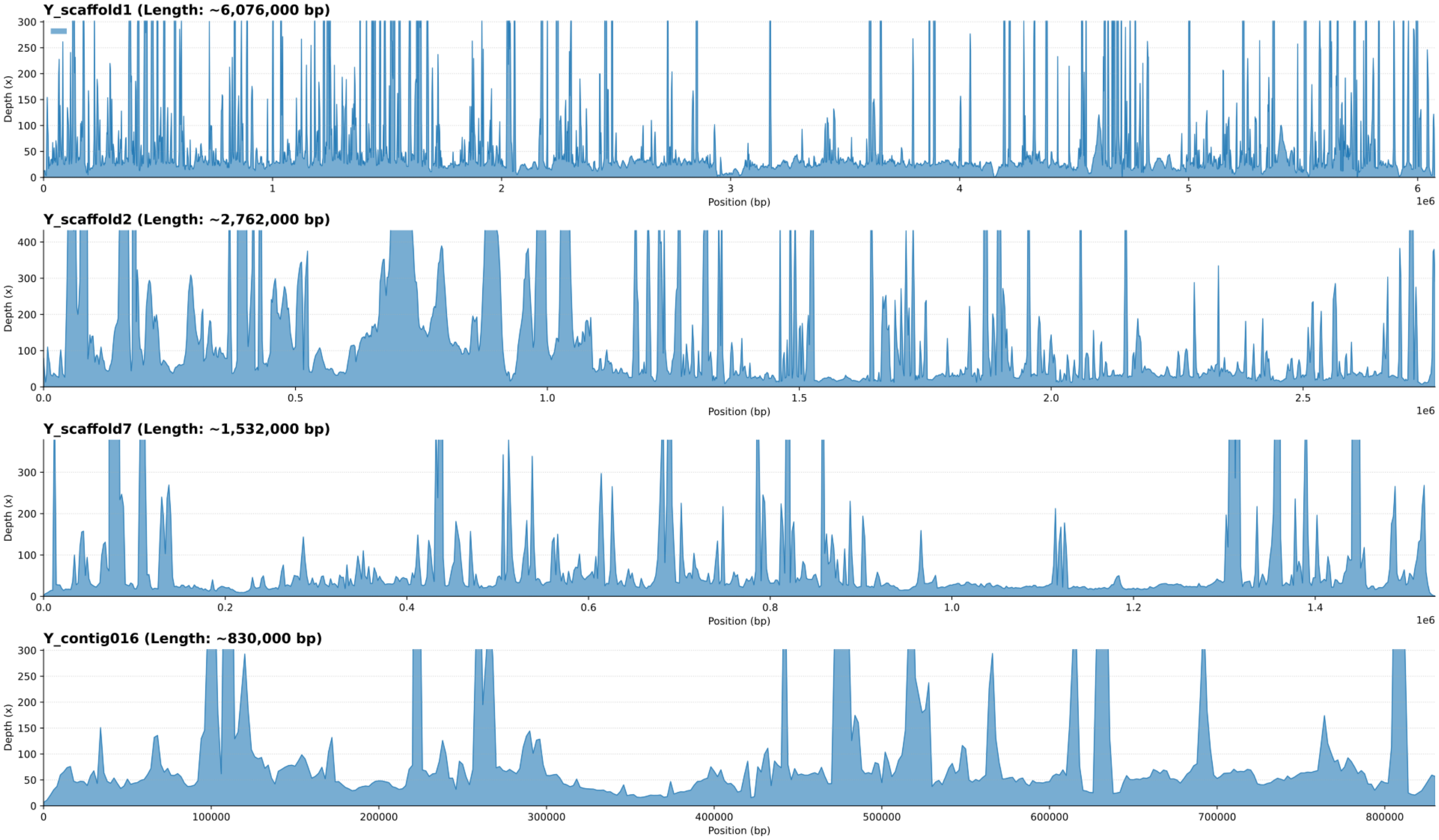

Supplementary Figure S2. PacBio HiFi read depth across scaffolded A3 Y-linked contigs. Coverage is broadly uniform across contigs, with localized spikes reflecting repetitive regions.

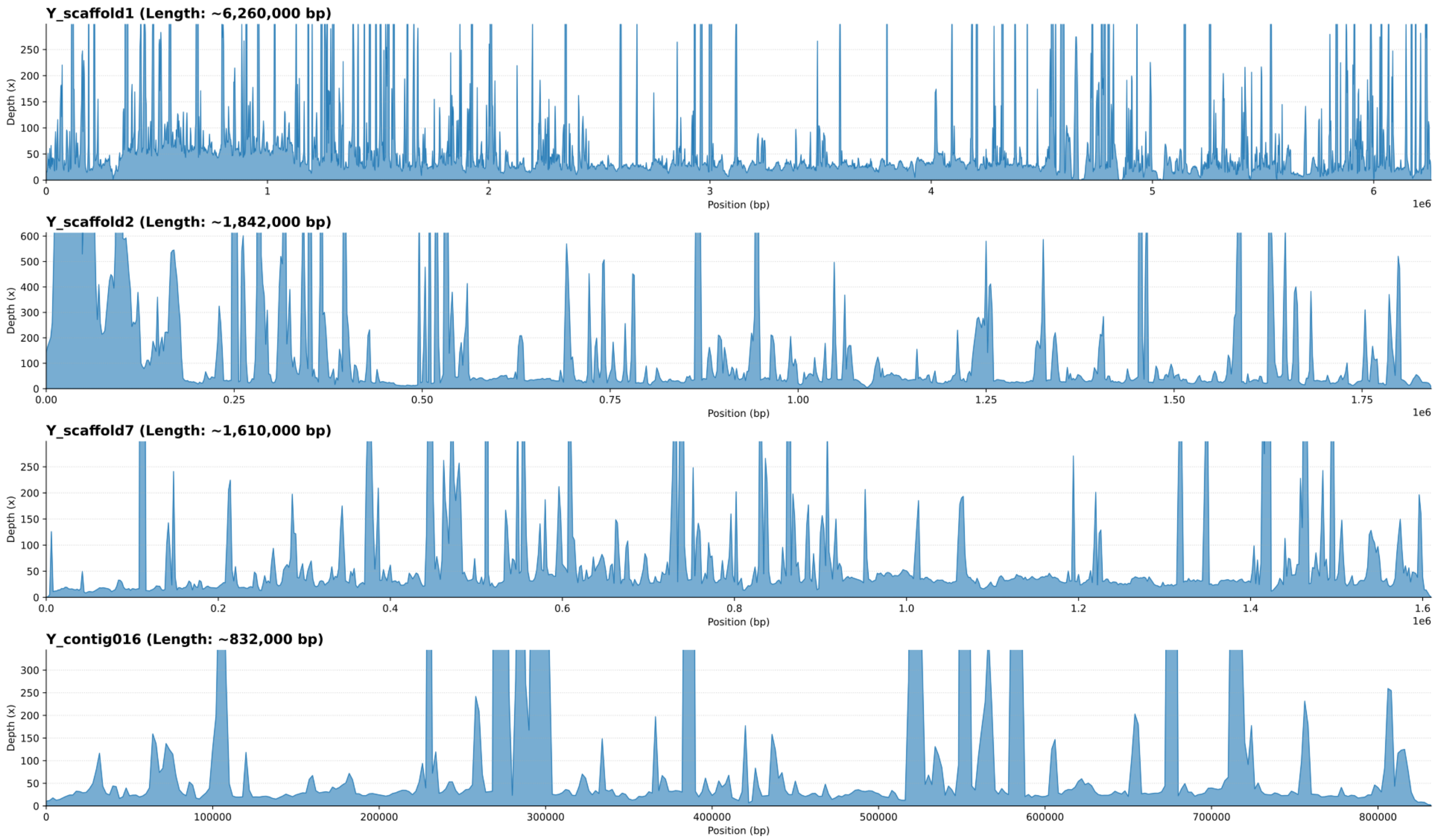

Supplementary Figure S3. PacBio HiFi read depth across scaffolded A4 Y-linked contigs. Coverage patterns are consistent with those observed in A3, supporting the continuity of assembly across the scaffolded Y-linked contigs.

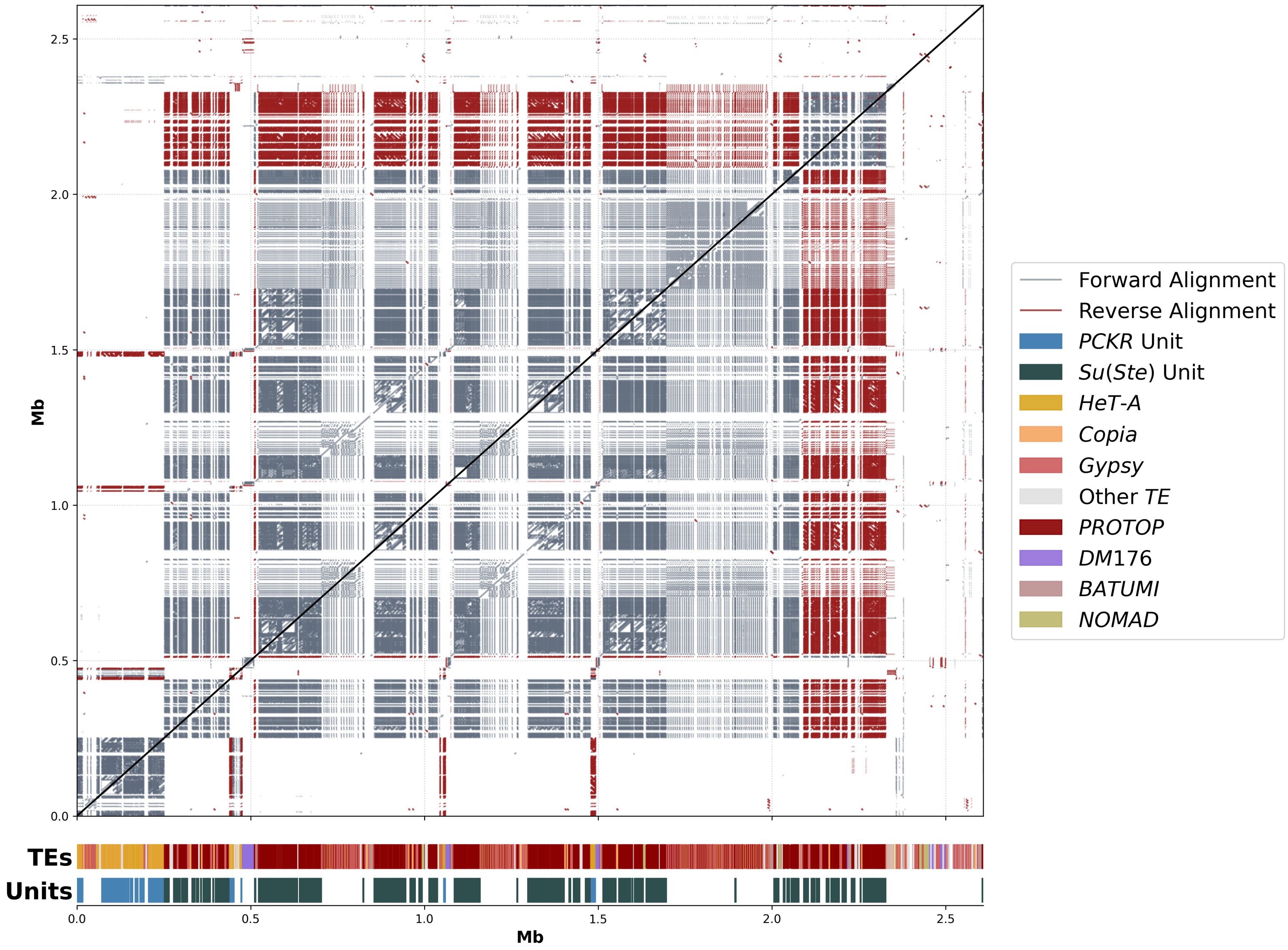

Supplementary Figure S4. Self-alignment MUMmer dot plot of the iso-1 *PCKR/Su(Ste)* array. Internal block structure and repeated-sequence organization are visible in off-diagonal alignments, reflecting sequence similarity among repeat units and higher-order array organization.

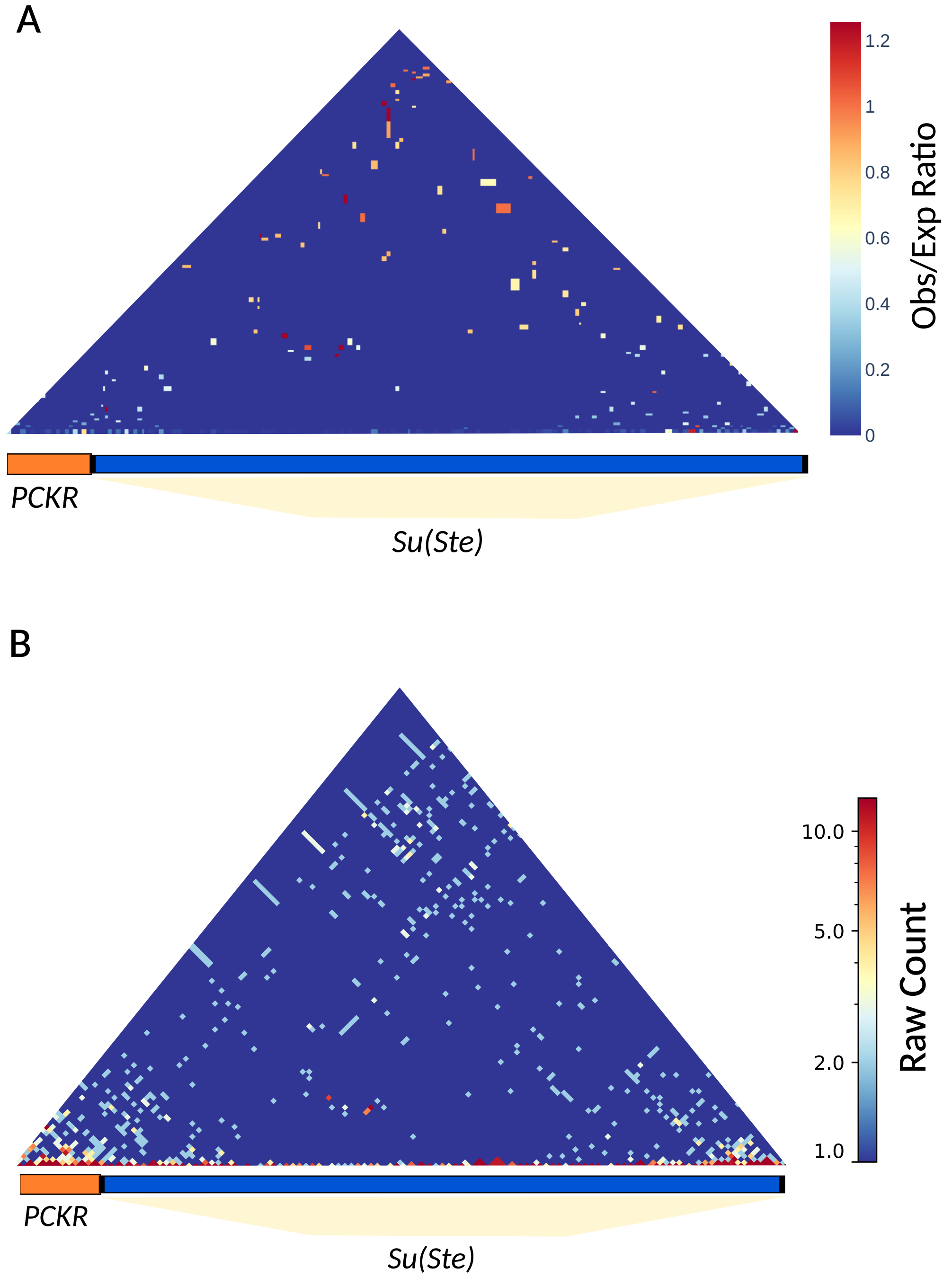

Supplementary Figure S5. Hi-C contact map of the iso-1 *Su(Ste)* and *PCKR* array. Contact maps are shown as observed/expected ratios (Panel A) and raw contact counts (Panel B)(Schauer et al. 2017). The interaction signal is generally low across the array, reflecting the repetitive and heterochromatic nature of this region.

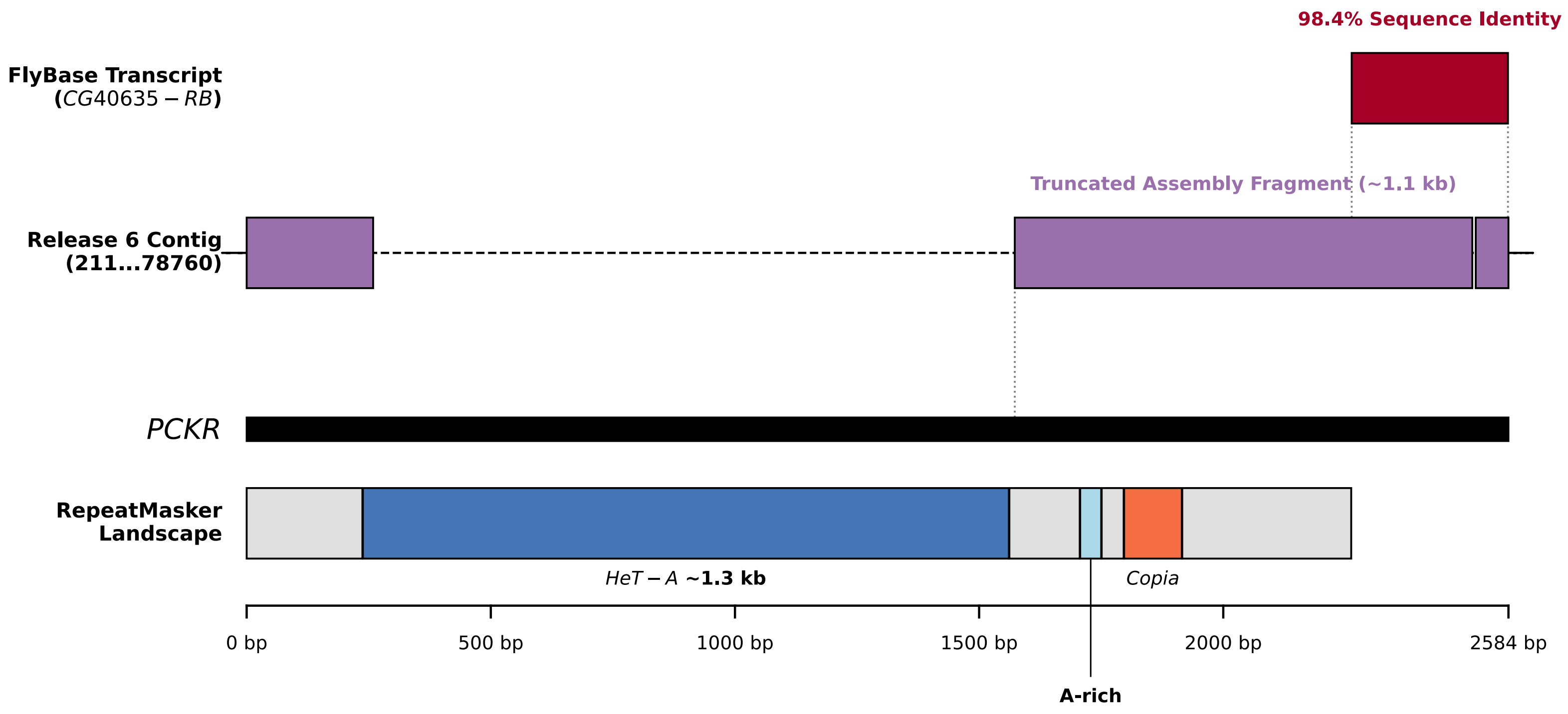

Supplementary Figure S6. Annotation of a *PCKR* repeat unit. The iso-1 *PCKR* unit from the Y assembly is aligned to the FlyBase *CG40635* transcript and its corresponding Release 6 contig, illustrating conservation of coding sequence alongside TE insertions.

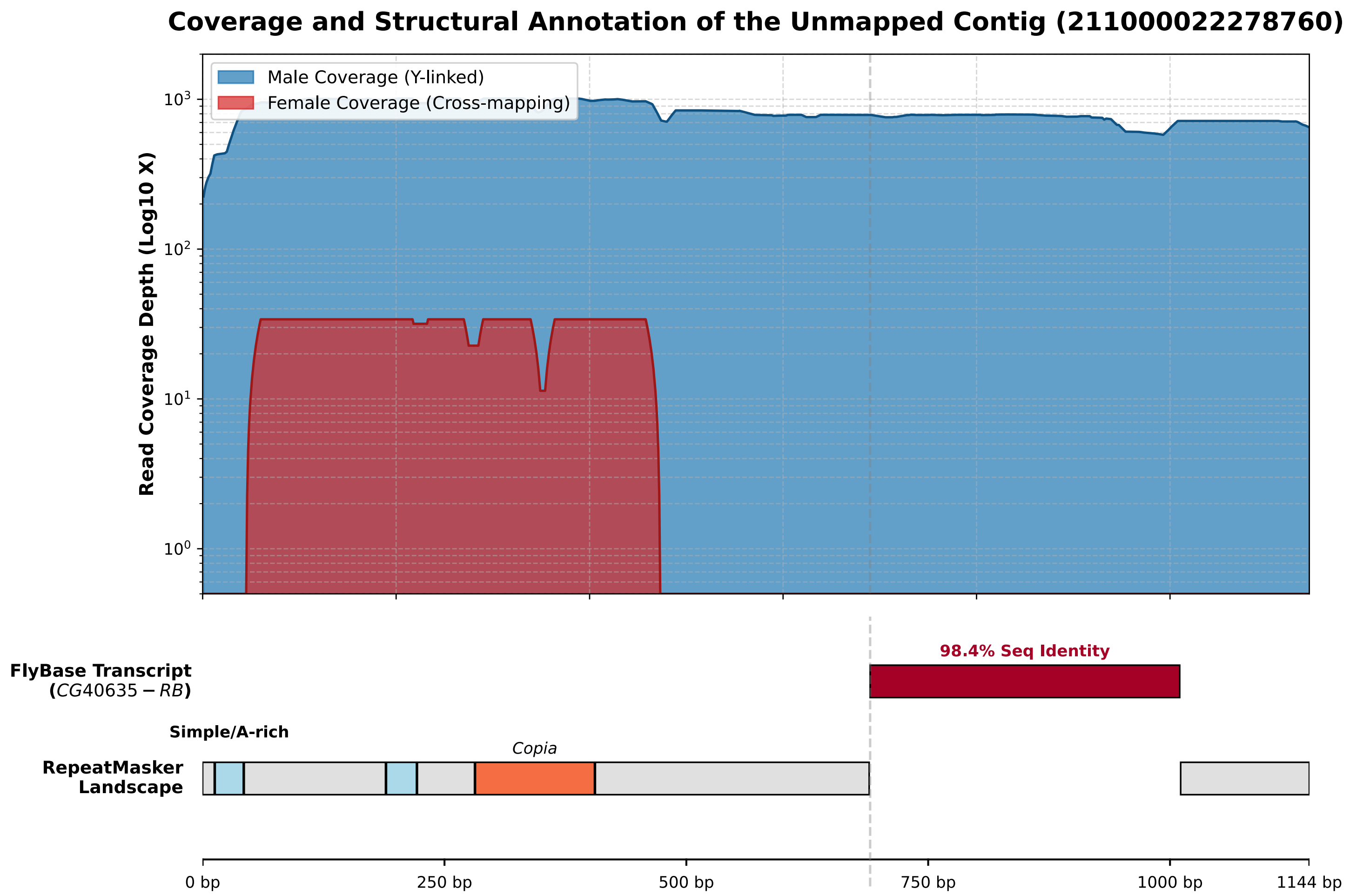

Supplementary Figure S7. Mapping of male and female A4 Nanopore direct RNA-seq reads to a Y-linked contig containing *CG40635*. Male-specific expression is evident from strong Y-linked coverage, while the female signal reflects cross-mapping due to high sequence similarity.

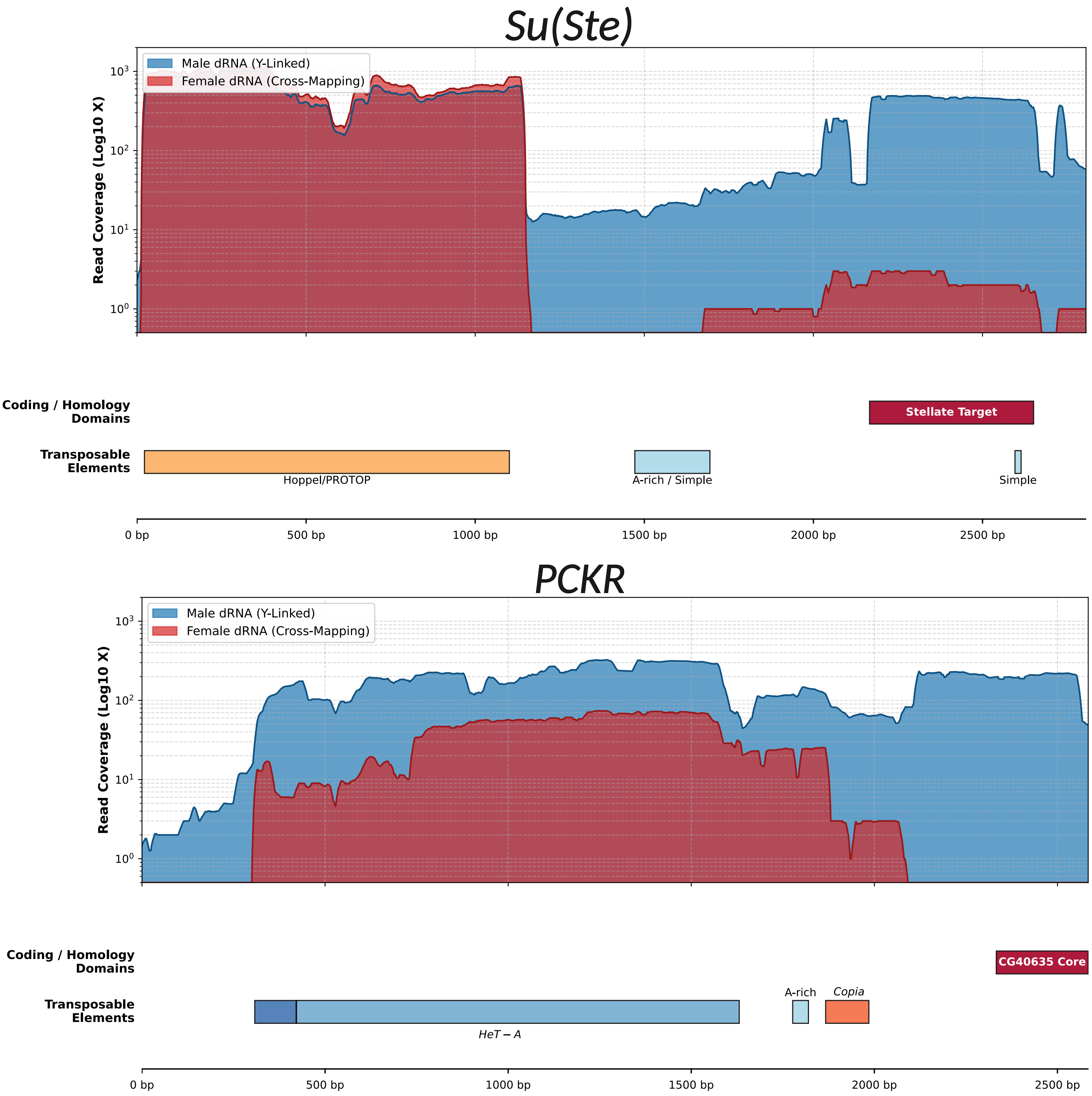

Supplementary Figure S8. Mapping of Nanopore RNA-seq reads to a representative *Su(Ste)* unit. Expression is predominantly male-derived, consistent with Y-linked transcription. Mapping of Nanopore RNA-seq reads to a representative *PCKR* unit. Greater coverage in male samples reflects Y-linked transcription, with the female coverage attributable to cross-mapping.

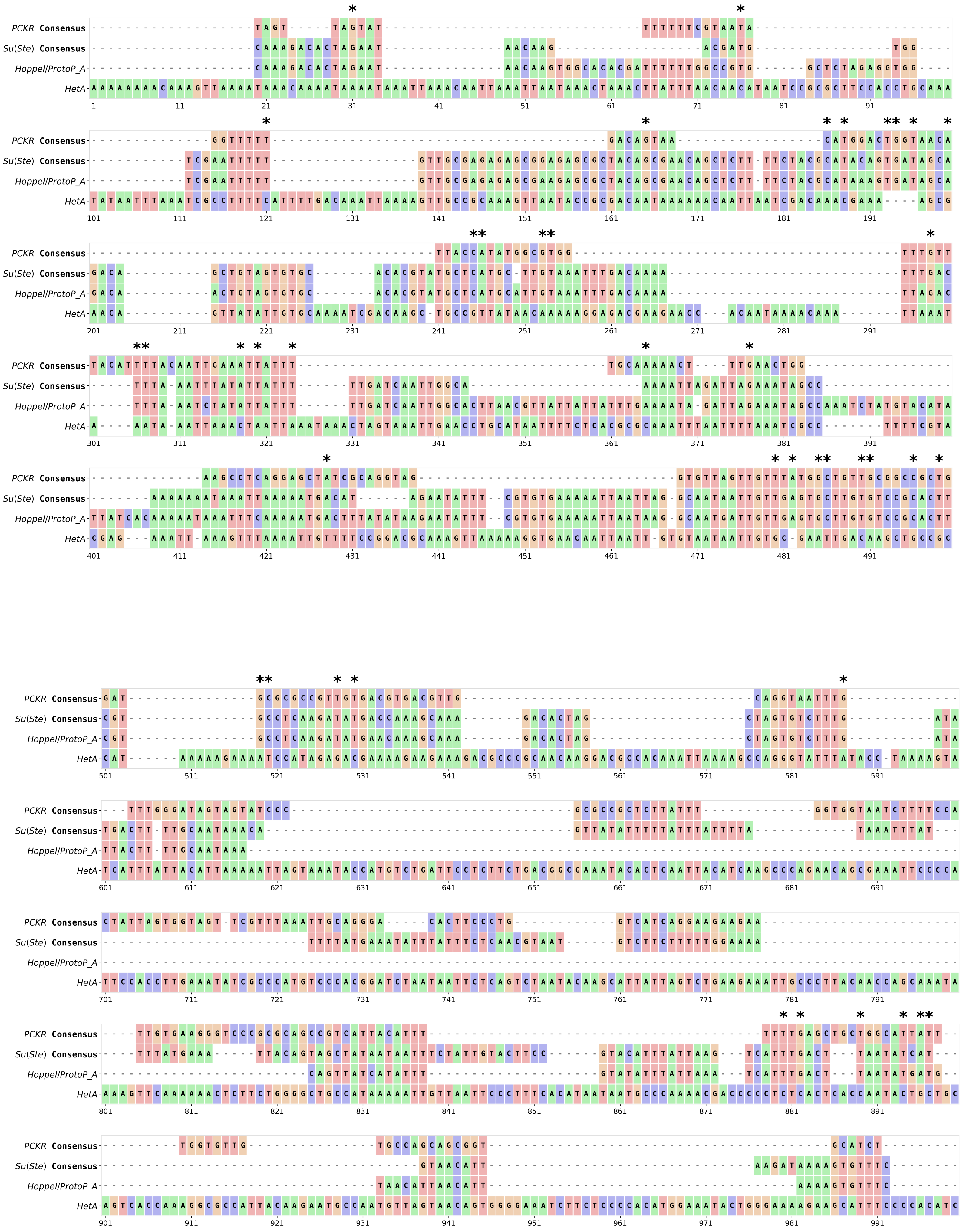
PCKR Consensus
Su(Ste) Consensus
Hoppel/ProtoP_A
HetA

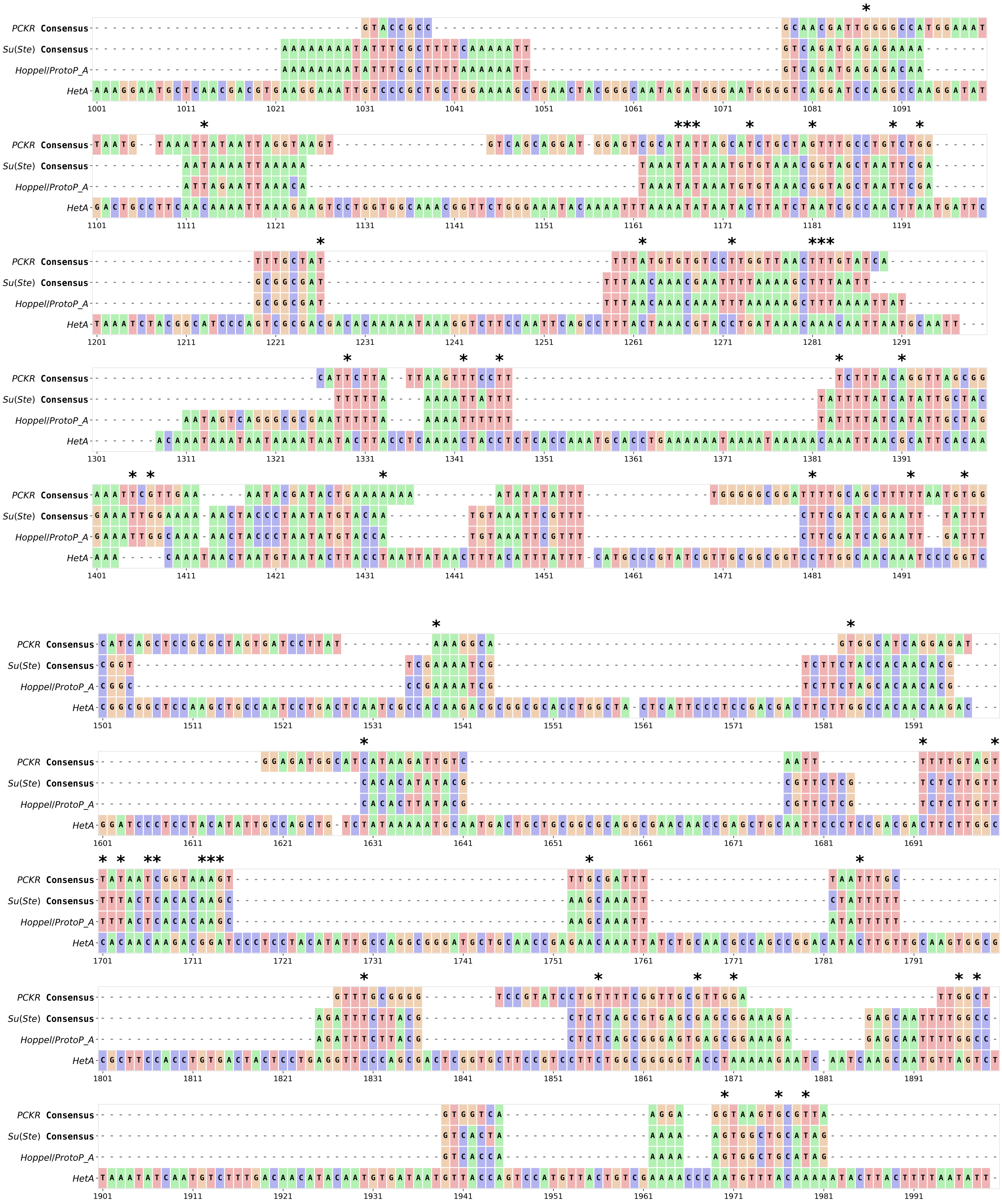
PCKR Consensus
Su(Ste) Consensus
Hoppel/ProtoP_A
HetA
1001
1101
1201
1301
1401
1501
1601
1701
1801
1901

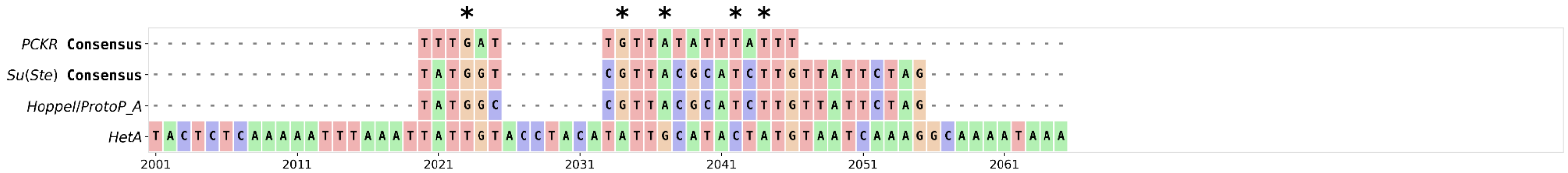

Supplementary Figure S9. Multiple sequence alignment of representative *Su(Ste)* and *PCKR* repeating units from the ISO-1 assembly, aligned against the canonical consensus sequences for *HetA* and *Hoppel* elements from a custom repeat library. Asterisks (*) placed above individual nucleotide positions highlight shared nucleotides present in both the *PCKR* and *Su(Ste)* sequences, but absent from the reference *HetA* sequence. Each nucleotide position is uniquely color-coded by base identity

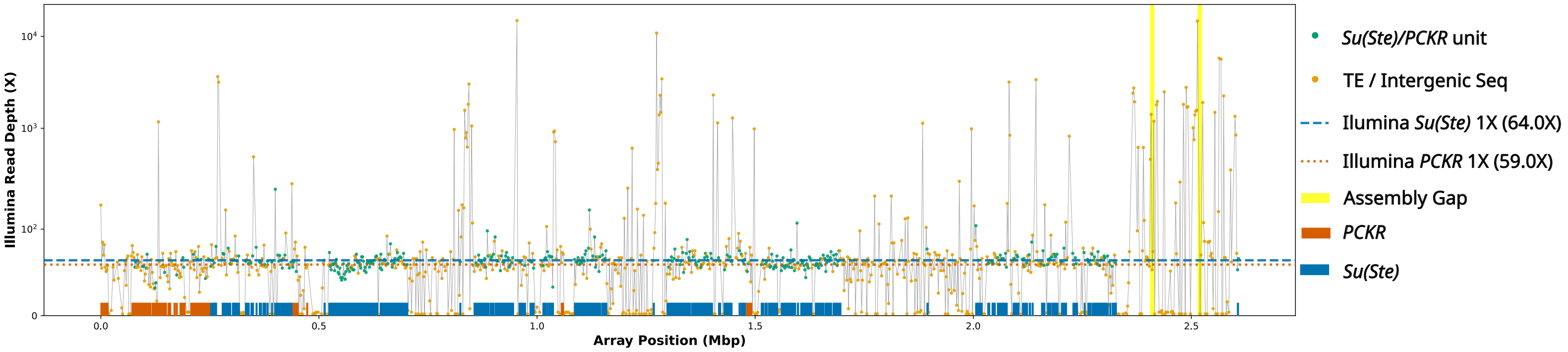

Supplementary Figure S10. Illumina short-read coverage across the merged *PCKR/Su(Ste)* array. Reads were mapped to a repeat-masked reference to assess coverage across the array. Dashed lines indicate estimated single-copy coverage baselines for *Su(Ste)* (red) and *PCKR* (blue), supporting assembly continuity.

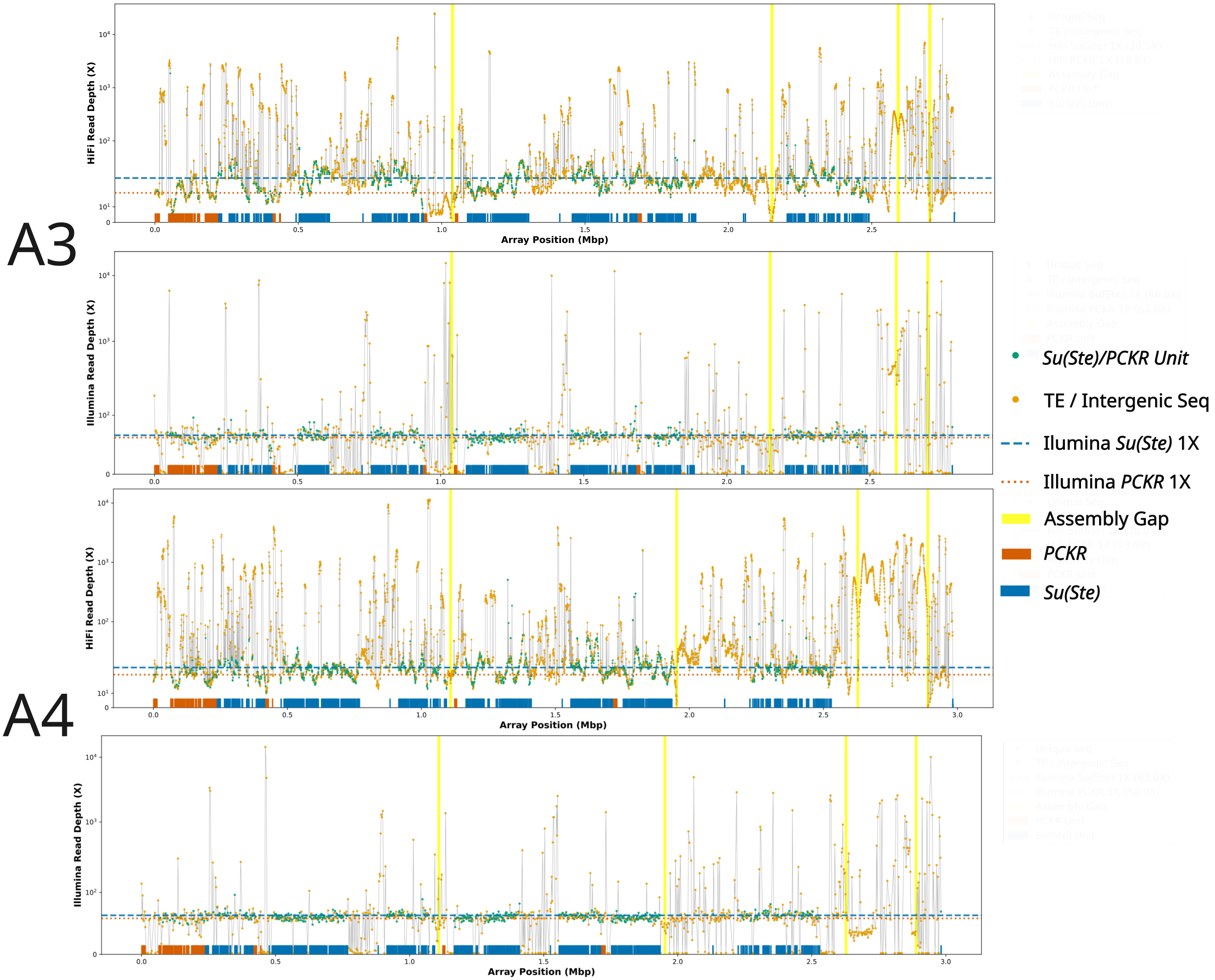

Supplementary Figure S11. Read depth across P*CKR/Su(Ste)* arrays in A3 and A4. HiFi (top panels) and Illumina (bottom panels) coverage profiles show uniform depth across the *Su(Ste)* and *PCKR* sequences. Dashed lines indicate estimated coverage baselines, supporting accurate representation of repeat copy numbers.

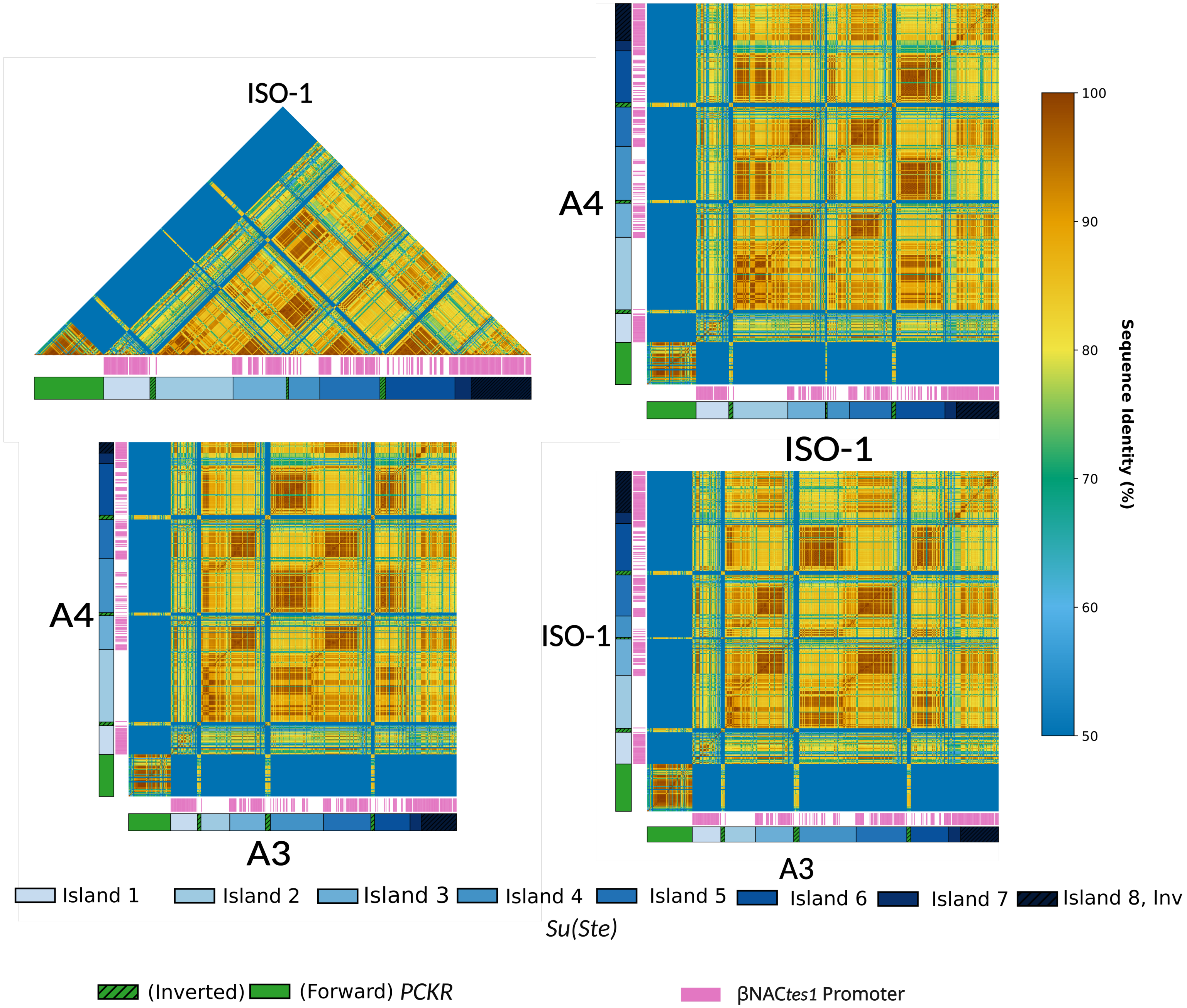

Supplementary Figure 12. Gap-penalized sequence similarity heatmaps for *PCKR* and *Su(Ste)* repeat units across iso-1, A3, and A4. Although overall island structure is retained, similarity patterns are reduced in TE-rich regions under gap-penalizing alignment, highlighting the influence of TE insertions on sequence similarity measurements.

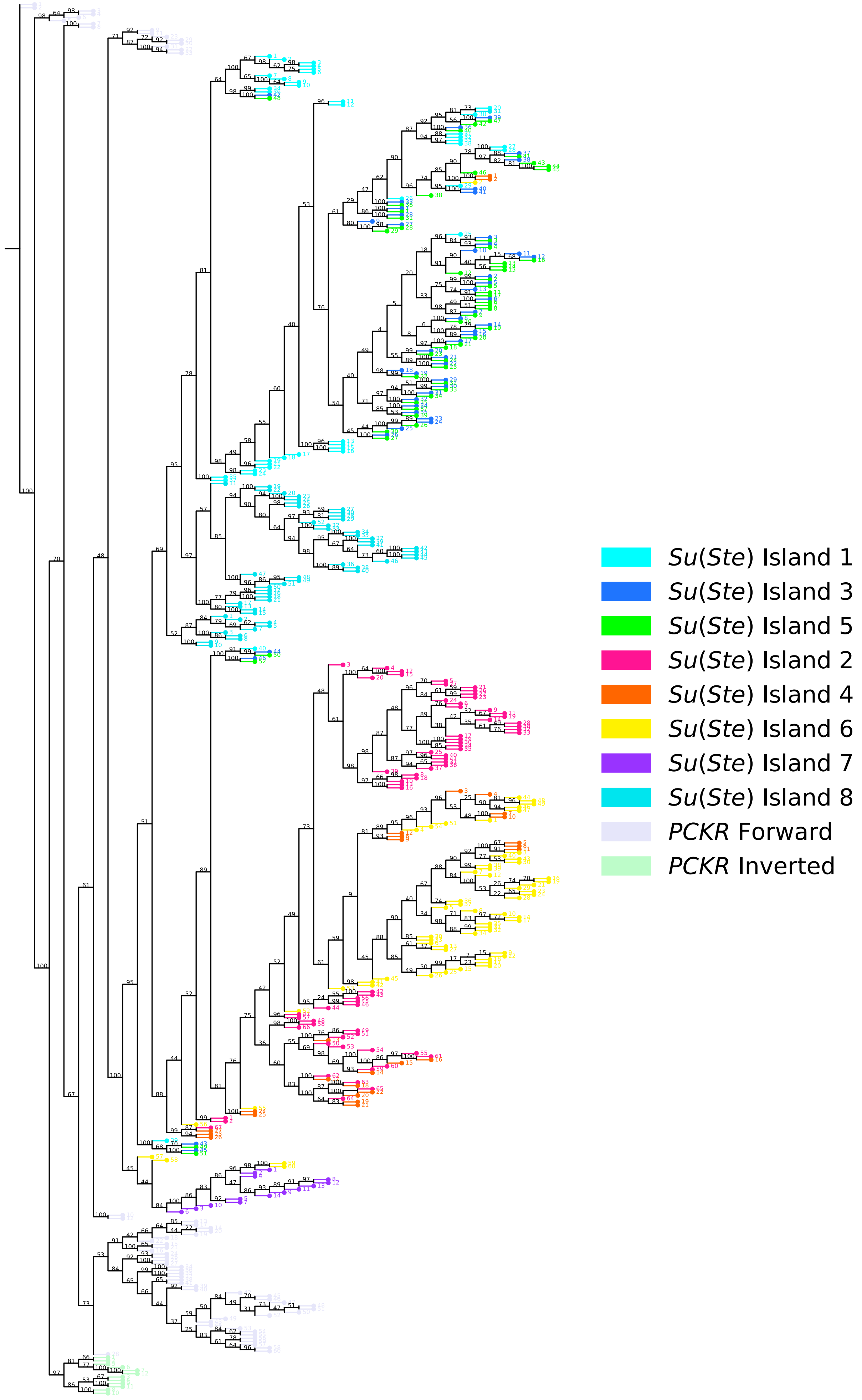

Supplementary Figure 13. Phylogenetic relationships among *Su(Ste)* and *PCKR* repeat units. Units cluster according to their position within the eight-island architecture, consistent with domain-restricted homogenization.

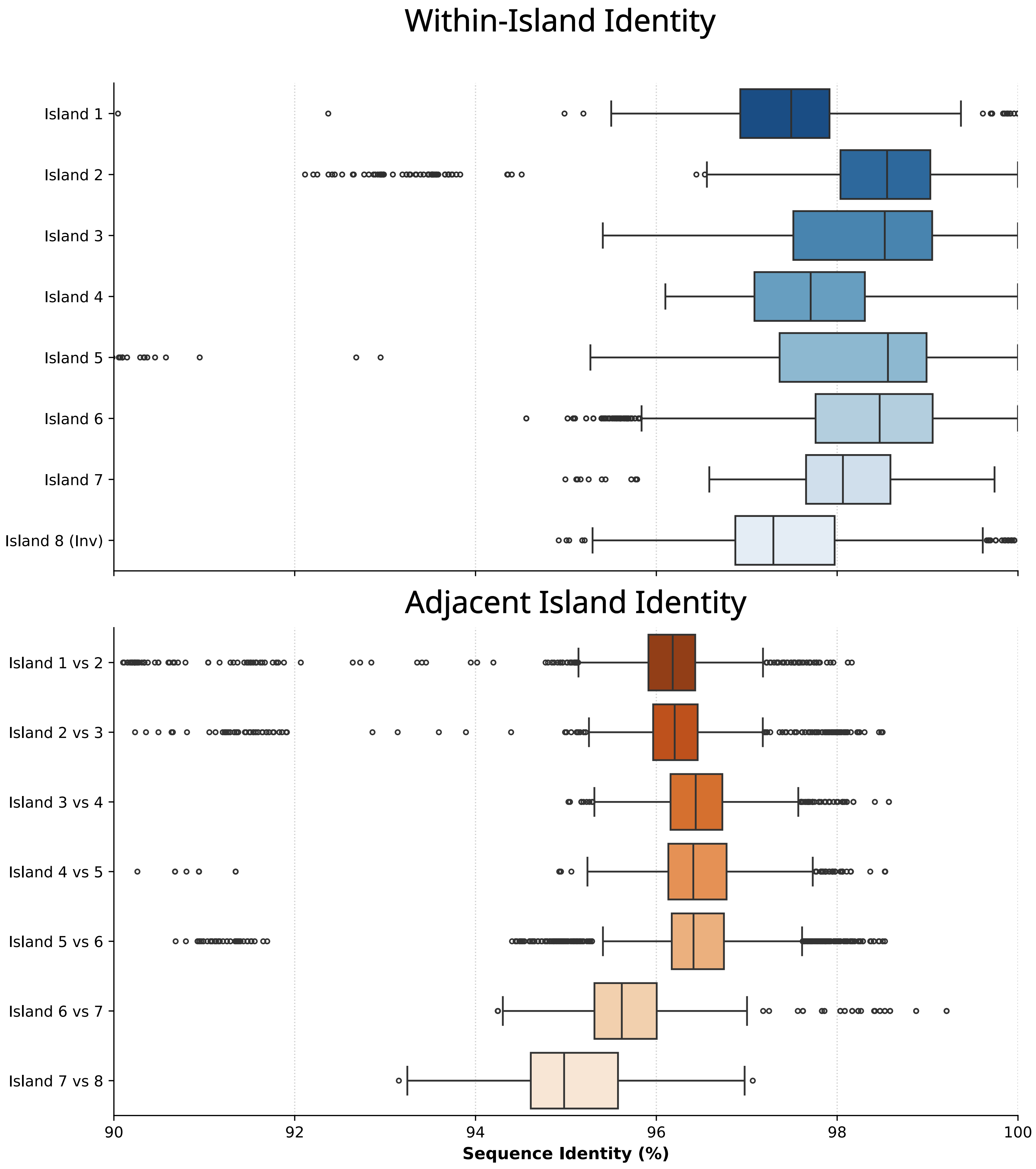

Supplementary Figure 14. Distributions of sequence identity within and between *Su(Ste)* islands. Higher sequence identity is observed within islands relative to adjacent islands, consistent with localized homogenization.

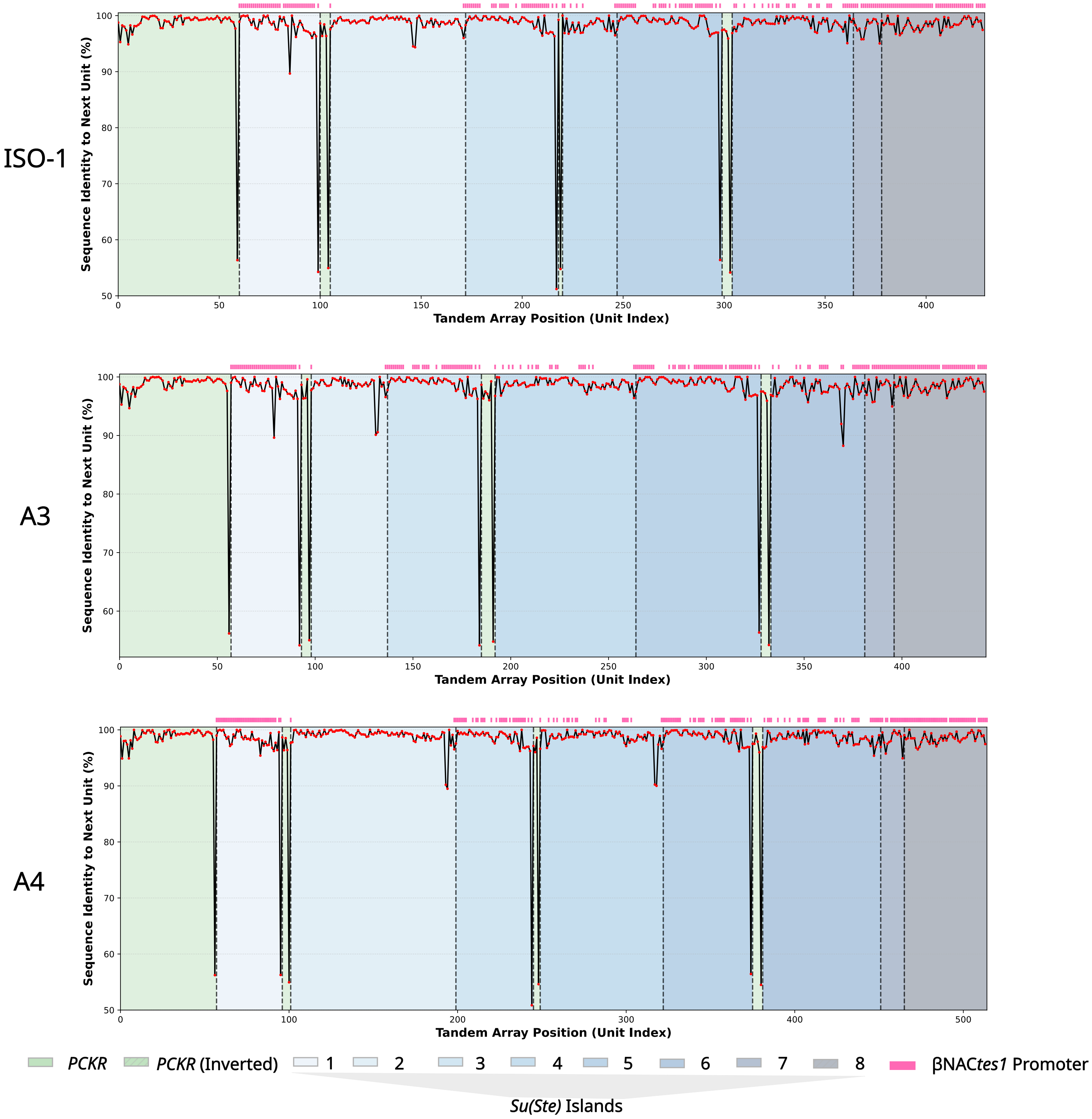

Supplementary Figure 15. Sequence identity between adjacent repeat units across *PCKR/Su(Ste)* arrays. Sharp reductions in identity occur at island boundaries, coinciding with transitions between repeat domains. *β*NAC*tes1* promoter locations are indicated.

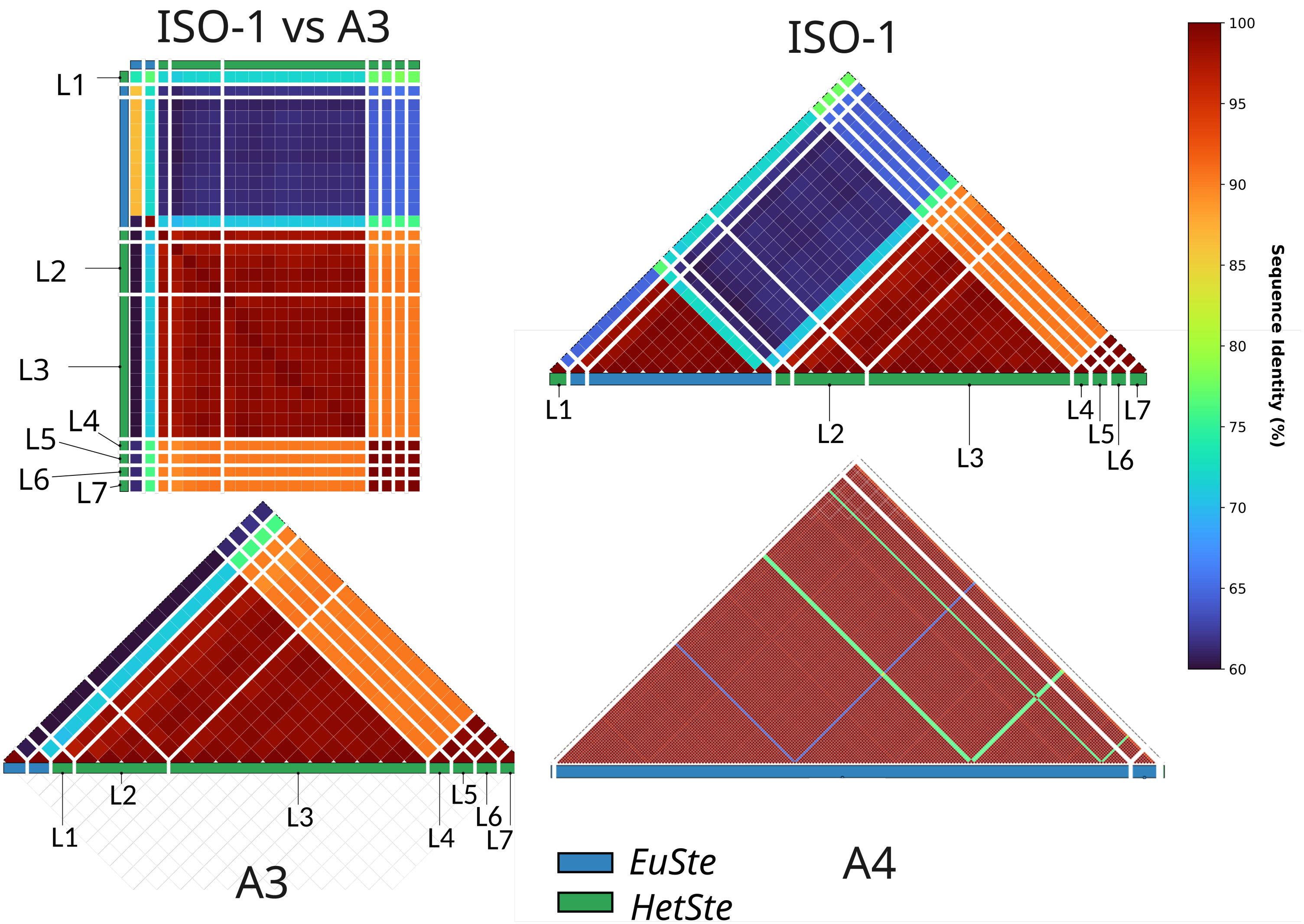

Supplementary Figure 16. Sequence similarity among X-linked *Stellate* repeat units. Heatmaps show pairwise identity within euchromatic and heterochromatic *Stellate* domains, revealing distinct patterns of homogenization relative to Y-linked *Su(Ste)* arrays. The white strips indicate physical separation between *Su(Ste)* sequences.

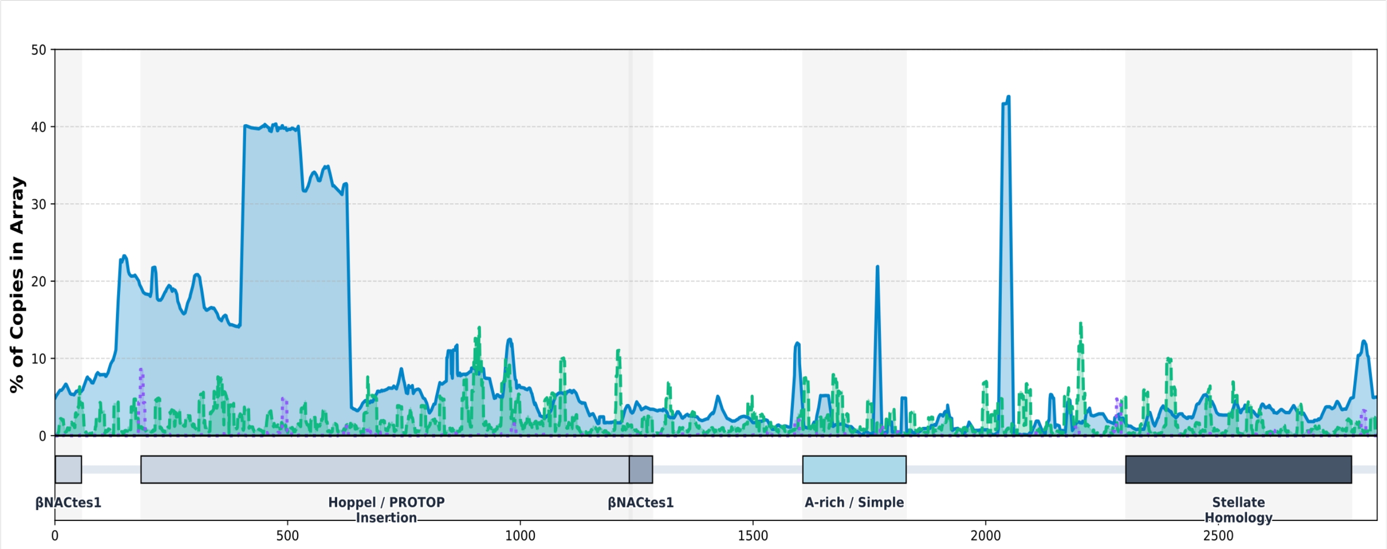
% of Copies in Array
βNACtes1
Hoppel / PROTOP Insertion
βNACtes1
A-rich / Simple
Stellate Homology

ISO-1

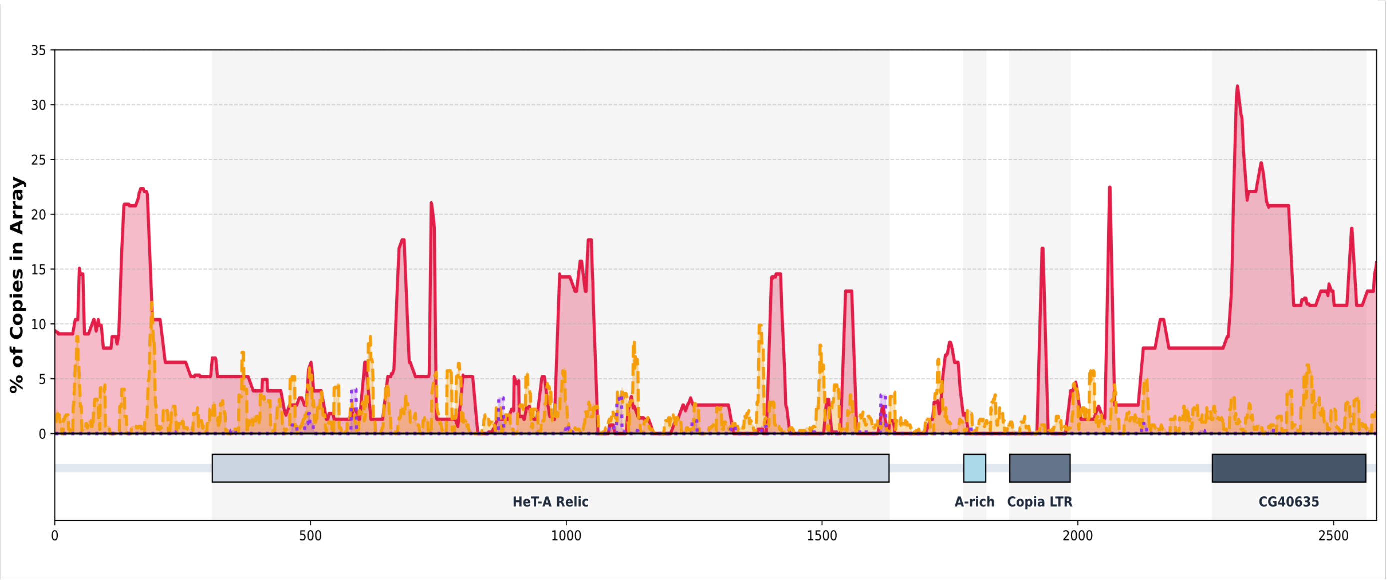
% of Copies in Array
HeT-A Relic
A-rich
Copia LTR
CG40635

A3

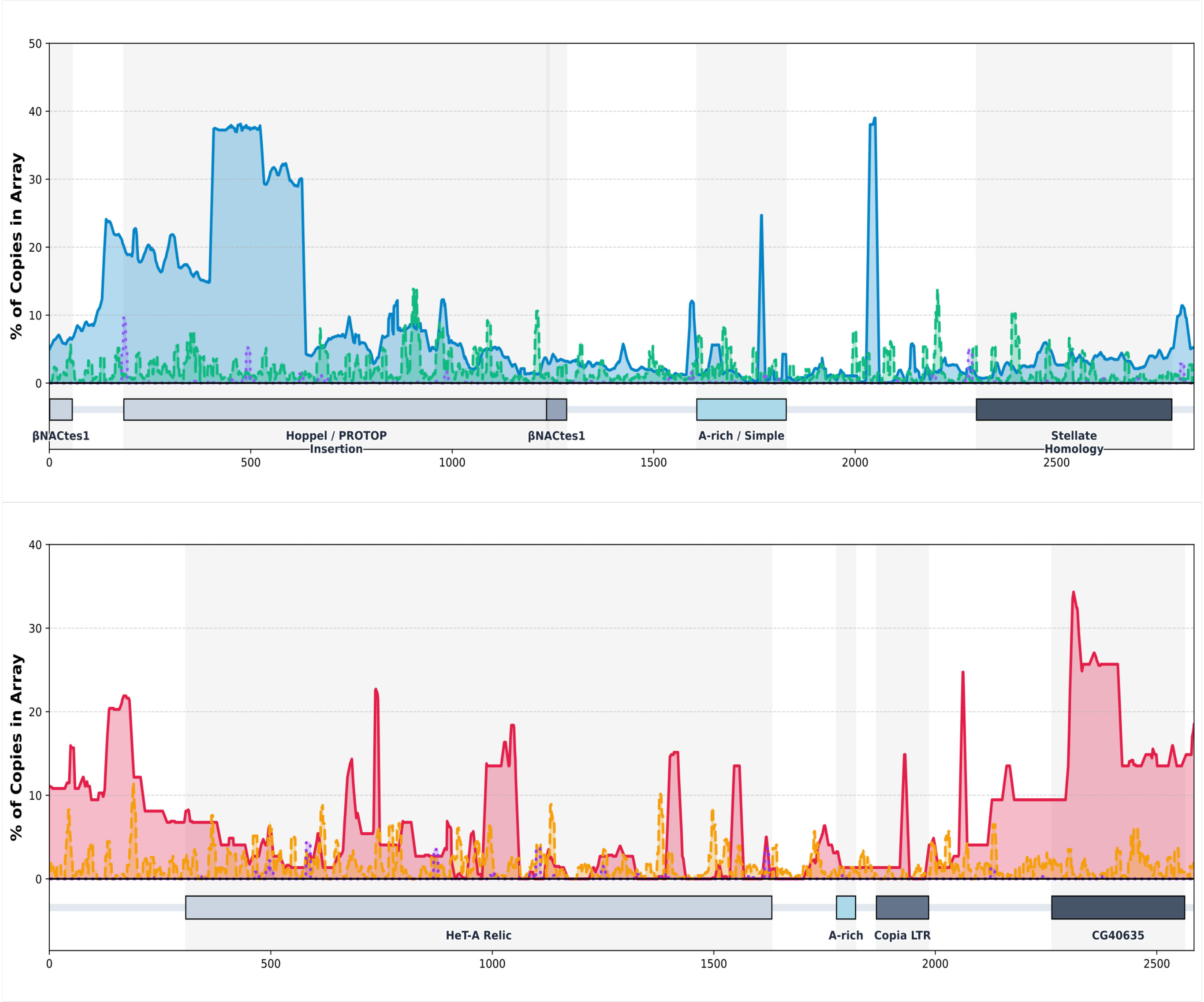
% of Copies in Array
βNACtes1
Hoppel / PROTOP Insertion
βNACtes1
A-rich / Simple
Stellate Homology
0
500
1000
1500
2000
2500
0
10
20
30
40
50
% of Copies in Array
HeT-A Relic
A-rich
Copia LTR
CG40635
0
500
1000
1500
2000
2500
0
10
20
30
40

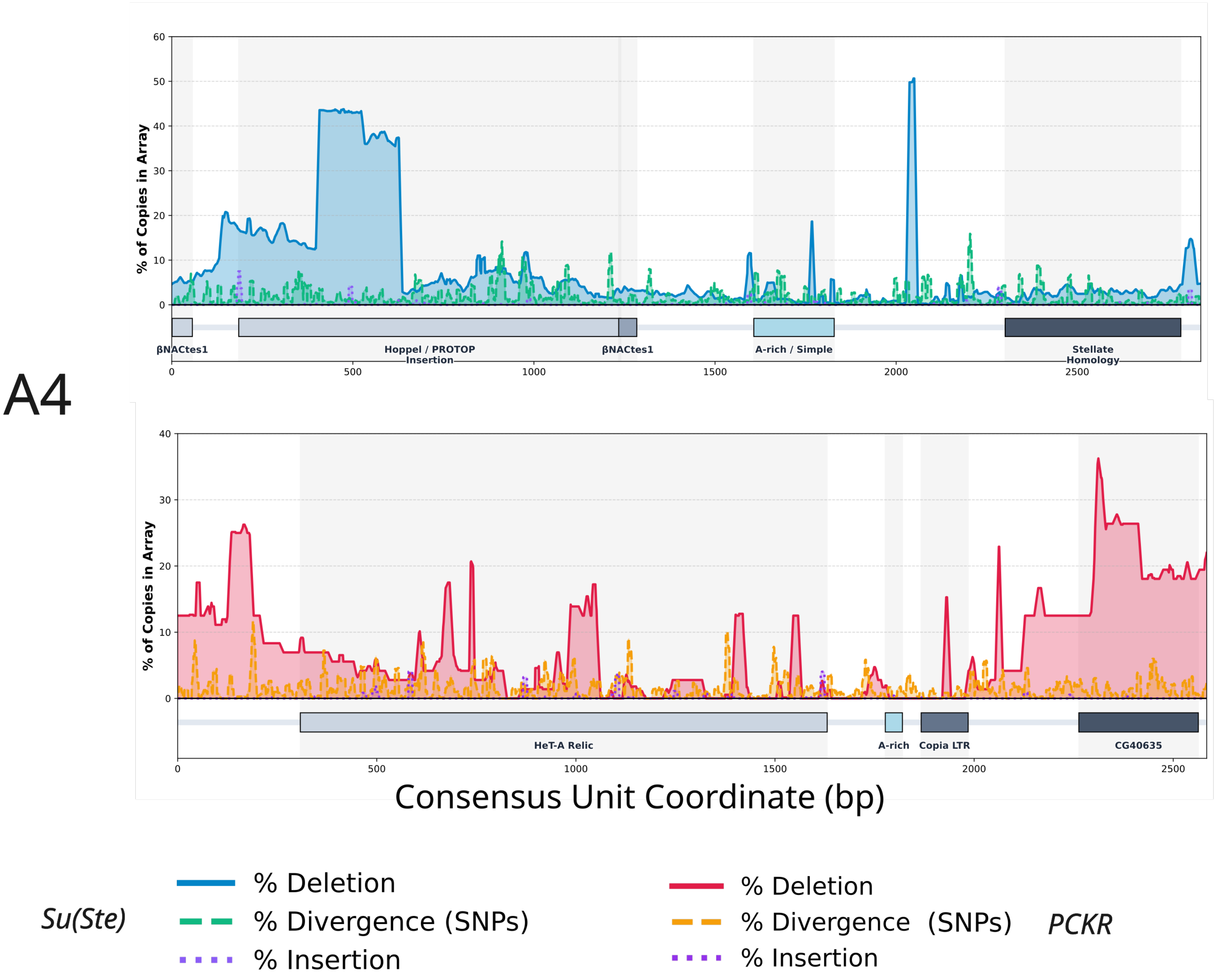

Supplementary Figure 17. Sequence variation across *PCKR* and *Su(Ste)* repeat units. Profiles of insertions, deletions, and SNPs show that variation is unevenly distributed along repeat units, with conserved regions forming a shared sequence backbone.

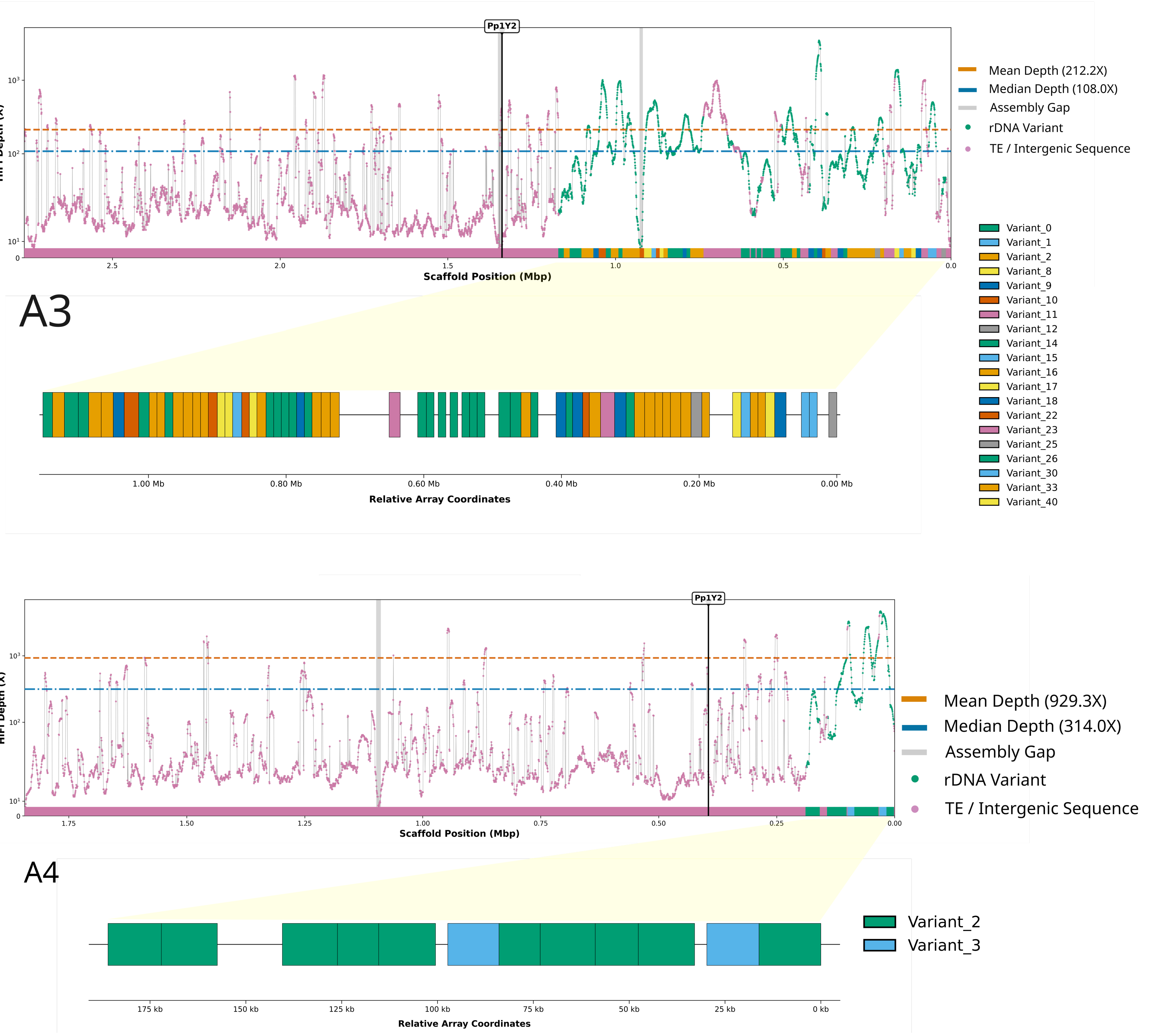

Supplementary Figure 18. Read mapping and RIBOTIN variant representation in A3 and A4 Y-linked rDNA arrays. Elevated coverage of HiFi read coverage suggests partial collapse of rDNA repeats. Mapping RIBOTIN-assembled rDNA variants to each array shows that the assemblies capture a subset of RIBOTIN-derived variants across arrays.

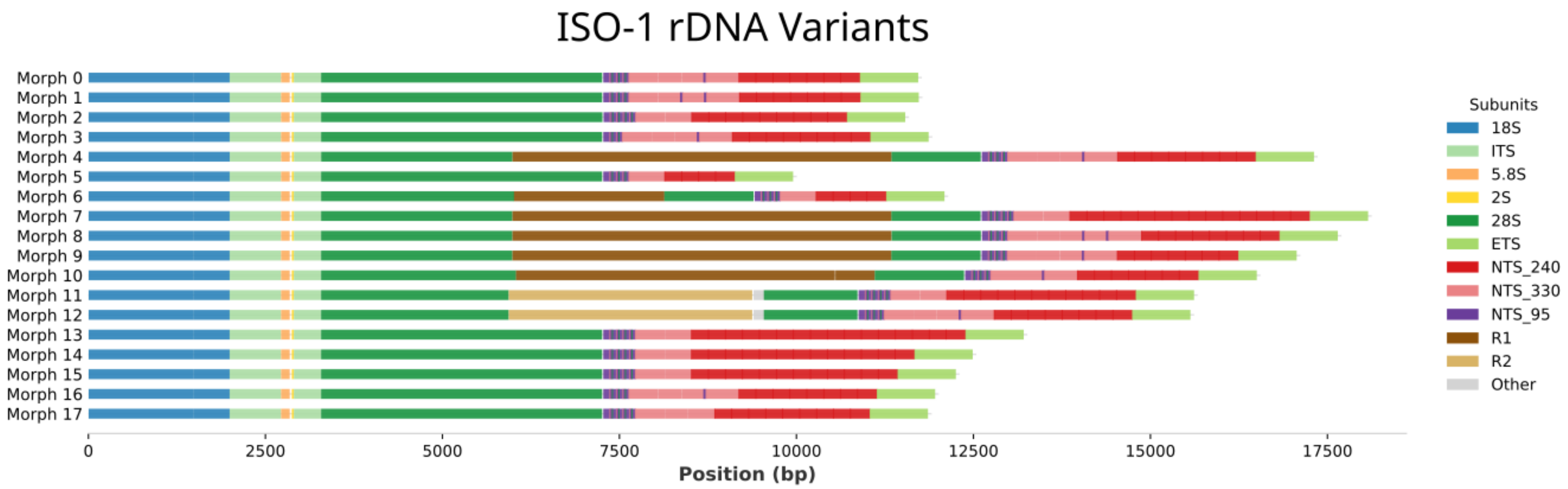

Supplementary Figure 19. RIBOTIN-derived rDNA variants from the iso-1 Y-linked array (see Methods). Variant structures show conserved coding regions alongside variable intergenic segments, reflecting heterogeneity within the array.

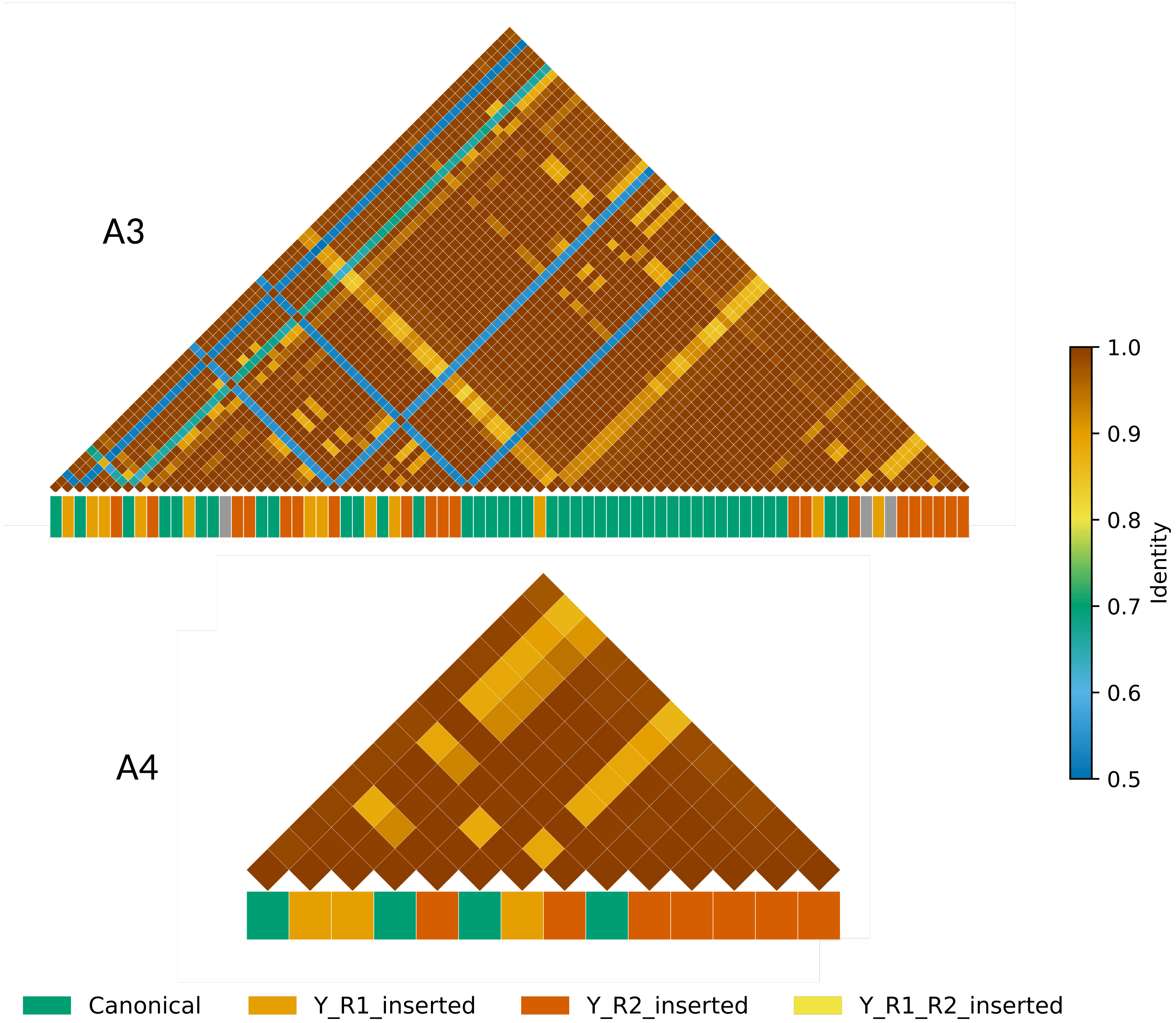

Supplementary Figure 20. Pairwise sequence alignment heatmaps showcasing sequence identity within the Y-linked rDNA units in the A3 and A4 assemblies. Annotated based on the *R1* and *R2* transposon insertion status.

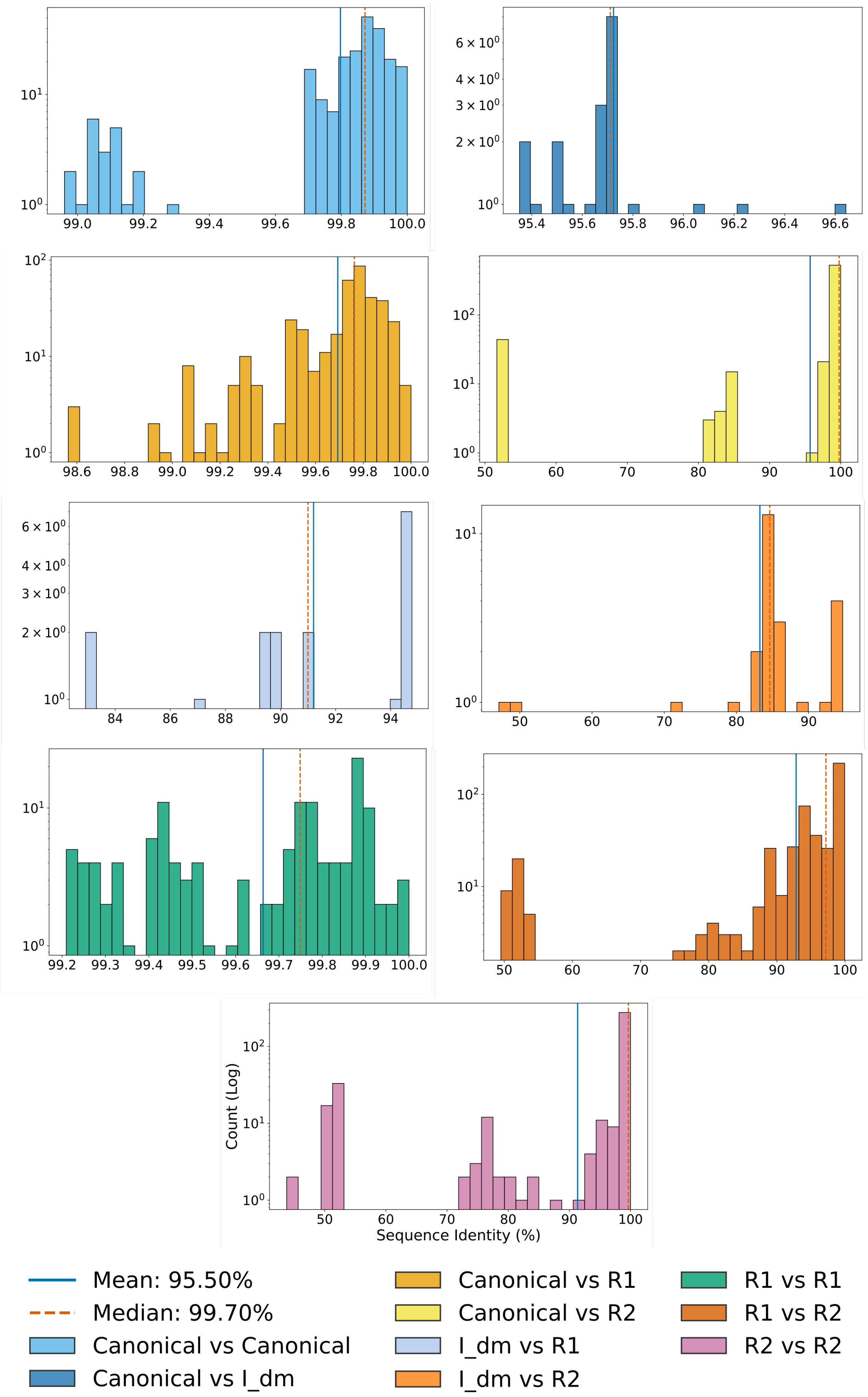

Supplementary Figure 21. Histograms show the pairwise distribution of sequence identity across different categories of rDNA repeat variants in the iso-1 Y chromosome assembly. Individual plots show specific alignment pairs, categorized by transposon insertion type (*R1, R2,* or *I-element*). "Canonical" units represent units without any insertions. The mean and Median in the figure legend are obtained by averaging the values from each individual plot.

Supplemental Table 1. Oxford Nanopore direct RNA-sequencing reads from A4 males and females, and BL156 males mapped to the iso-1 assembly reveal transcript support for the major fertility gene annotations in our assembly. Annotations were validated via comparison to the Chang et al. (2019) heterochromatin-enriched assembly. Cross-mapping of A4 female transcripts to the *FDY* gene is due to the very high sequence identity between *FDY* and its autosomal counterpart, *vig2.*

| Column 1 | Exon | Scaffold | Start | End | Length | A4_Male | A4_Female | BL156_Male |
|---|---|---|---|---|---|---|---|---|
| *ARY* | 1 | Y_Scaffold1 | 2773910 | 2774029 | 120 | 136.46 | 0 | 7.3 |
| *ARY* | 2 | Y_Scaffold1 | 2773629 | 2773796 | 168 | 139.01 | 0 | 9.83 |
| *ARY* | 3 | Y_Scaffold1 | 2773408 | 2773577 | 170 | 148.64 | 0 | 12 |
| *ARY* | 4 | Y_Scaffold1 | 2773004 | 2773346 | 343 | 153.68 | 0 | 14.81 |
| *ARY* | 5 | Y_Scaffold1 | 2772648 | 2772947 | 300 | 176.96 | 0 | 19.4 |
| *CCY* | 1 | Y_scaffold5 | 28287 | 29357 | 1071 | 41.4 | 0 | 0.9 |
| *CCY* | 2 | Y_scaffold5 | 49389 | 49544 | 156 | 34.56 | 0 | 0.95 |
| *CCY* | 3 | Y_scaffold5 | 222515 | 225167 | 2653 | 39.95 | 0 | 1.85 |
| *CCY* | 4 | Y_scaffold5 | 225223 | 225344 | 122 | 45.05 | 0 | 2.56 |
| *CG41561* | 1 | Y_Contig74 | 14212 | 14255 | 44 | 112.8 | 0 | 11.98 |
| *CG41561* | 2 | Y_Contig74 | 14310 | 14536 | 227 | 113.88 | 0 | 14.44 |
| *CG41561* | 3 | Y_Contig74 | 15139 | 16108 | 970 | 130.49 | 0 | 17.86 |
| *CG41561* | 4 | Y_Contig74 | 16167 | 16245 | 79 | 144.28 | 0 | 20.32 |
| *FDY* | 1 | Y_Contig1 | 257757 | 258228 | 472 | 608.76 | 8220.93 | 200.97 |
| *FDY* | 2 | Y_Contig1 | 256768 | 257285 | 518 | 718.53 | 9166.03 | 252.93 |
| *ORY* | 1 | Y_scaffold4 | 714829 | 714990 | 162 | 1.95 | 0 | 0.84 |
| *ORY* | 2 | Y_scaffold4 | 714412 | 714761 | 350 | 1.92 | 0 | 0.85 |
| *ORY* | 3 | Y_scaffold4 | 530414 | 531037 | 624 | 11.32 | 0 | 2.13 |
| *ORY* | 4 | Y_scaffold4 | 530043 | 530359 | 317 | 12.73 | 0 | 3.07 |
| *ORY* | 5 | Y_scaffold4 | 406182 | 406360 | 179 | 13.66 | 0 | 3.68 |
| *ORY* | 6 | Y_scaffold4 | 405258 | 405901 | 644 | 15.01 | 0 | 4.35 |
| *ORY* | 7 | Y_scaffold4 | 405066 | 405204 | 139 | 17.31 | 0.21 | 5.9 |
| *ORY* | 8 | Y_scaffold4 | 141484 | 141813 | 330 | 19.64 | 0 | 5.38 |
| *PRY* | 1 | Y_Scaffold1 | 6509513 | 6510123 | 611 | 6.03 | 0 | 3.01 |
| *PRY* | 2 | Y_Scaffold1 | 6542101 | 6542758 | 658 | 8.59 | 0 | 2.41 |
| *PRY* | 3 | Y_Scaffold1 | 6572057 | 6574531 | 2475 | 4.31 | 0.01 | 0.22 |

| | | | | | | | | |
|---|---|---|---|---|---|---|---|---|
| *Pp1-Y1* | 1 | Y_Contig2 | 15178 | 16098 | 921 | 47.28 | 0 | 0 |
| *Pp1-Y2* | 1 | Y_Scaffold2 | 1239696 | 1240625 | 930 | 50.81 | 0 | 3.26 |
| *Ppr-Y* | 1 | Y_Scaffold1 | 1071316 | 1071442 | 127 | 0 | 0 | 0 |
| *Ppr-Y* | 2 | Y_Scaffold1 | 1422885 | 1423230 | 346 | 21.32 | 0 | 1.4 |
| *Ppr-Y* | 3 | Y_Scaffold1 | 1762271 | 1762847 | 577 | 28.35 | 0 | 0.92 |
| *Ppr-Y* | 4 | Y_Scaffold1 | 1818505 | 1818801 | 297 | 12.87 | 0 | 1.63 |
| *Ppr-Y* | 5 | Y_Scaffold1 | 1818803 | 1819083 | 281 | 16.15 | 0 | 1.51 |
| *Ppr-Y* | 6 | Y_Scaffold1 | 2378761 | 2378939 | 179 | 39.03 | 0 | 2.41 |
| *Ppr-Y* | 7 | Y_Scaffold1 | 2497575 | 2497858 | 284 | 48.85 | 0 | 1.88 |
| *Ppr-Y* | 8 | Y_Scaffold1 | 2601733 | 2601942 | 210 | 48.37 | 0 | 1.9 |
| *WDY* | 1 | Y_Scaffold1 | 280002 | 280377 | 376 | 23.91 | 0 | 0.7 |
| *WDY* | 2 | Y_Scaffold1 | 278030 | 279110 | 1081 | 24.67 | 0 | 0.83 |
| *WDY* | 3 | Y_Scaffold1 | 53698 | 54152 | 455 | 36.4 | 0 | 1.72 |
| *WDY* | 4 | Y_Scaffold1 | 53439 | 53637 | 199 | 37.34 | 0 | 1.65 |
| *WDY* | 5 | Y_Scaffold1 | 21128 | 21328 | 201 | 20 | 0 | 1.16 |
| *WDY* | 6 | Y_Scaffold1 | 20403 | 21066 | 664 | 24.35 | 0 | 1.67 |
| *kl-2* | 1 | Y_Scaffold1 | 5552807 | 5553473 | 667 | 6.35 | 0 | 0 |
| *kl-2* | 2 | Y_Scaffold1 | 5428856 | 5430705 | 1850 | 6.65 | 0 | 0 |
| *kl-2* | 3 | Y_Scaffold1 | 5321315 | 5321996 | 682 | 6.27 | 0 | 0 |
| *kl-2* | 4 | Y_Scaffold1 | 5320875 | 5321260 | 386 | 6.85 | 0 | 0 |
| *kl-2* | 5 | Y_Scaffold1 | 5276513 | 5276955 | 443 | 8.03 | 0 | 0 |
| *kl-2* | 6 | Y_Scaffold1 | 5254265 | 5257444 | 3180 | 10.64 | 0 | 0 |
| *kl-2* | 7 | Y_Scaffold1 | 5250127 | 5254208 | 4082 | 17.09 | 0 | 0 |
| *kl-2* | 8 | Y_Scaffold1 | 5249960 | 5250064 | 105 | 22.05 | 0 | 0 |
| *kl-2* | 9 | Y_Scaffold1 | 5248964 | 5249906 | 943 | 23.69 | 0 | 0.52 |
| *kl-2* | 10 | Y_Scaffold1 | 5248411 | 5248904 | 494 | 26.17 | 0 | 1.16 |
| *kl-2* | 11 | Y_Scaffold1 | 5247806 | 5248353 | 548 | 26.8 | 0 | 0.76 |
| *kl-3* | 1 | Y_Scaffold1 | 5770732 | 5771757 | 1026 | 10.2 | 0 | 0 |
| *kl-3* | 2 | Y_Scaffold1 | 5781972 | 5782075 | 104 | 11.86 | 0 | 0 |
| *kl-3* | 3 | Y_Scaffold1 | 5782132 | 5782307 | 176 | 11.85 | 0 | 0 |
| *kl-3* | 4 | Y_Scaffold1 | 5782362 | 5783473 | 1112 | 13.88 | 0 | 0.23 |
| *kl-3* | 5 | Y_Scaffold1 | 5832910 | 5833078 | 169 | 16.09 | 0 | 0.93 |
| *kl-3* | 6 | Y_Scaffold1 | 5833139 | 5833544 | 406 | 16.16 | 0 | 0.91 |

| | | | | | | | | |
|---|---|---|---|---|---|---|---|---|
| *kl-3* | 7 | Y_Scaffold1 | 5979953 | 5980082 | 130 | 12.15 | 0 | 0 |
| *kl-3* | 8 | Y_Scaffold1 | 5980136 | 5982348 | 2213 | 15.25 | 0.01 | 0.04 |
| *kl-3* | 9 | Y_Scaffold1 | 5982412 | 5983796 | 1385 | 21.4 | 0 | 0 |
| *kl-3* | 10 | Y_Scaffold1 | 6025793 | 6026127 | 335 | 20.53 | 0 | 0 |
| *kl-3* | 11 | Y_Scaffold1 | 6217877 | 6218685 | 809 | 21.95 | 0 | 0 |
| *kl-3* | 12 | Y_Scaffold1 | 6218960 | 6220420 | 1461 | 25.52 | 0 | 0 |
| *kl-3* | 13 | Y_Scaffold1 | 6291627 | 6294502 | 2876 | 32.81 | 0 | 0.33 |
| *kl-3* | 14 | Y_Scaffold1 | 6294558 | 6294736 | 179 | 39.64 | 0 | 1.34 |
| *kl-3* | 15 | Y_Scaffold1 | 6315240 | 6315911 | 672 | 42.58 | 0 | 2.07 |
| *kl-5* | 1 | Y_scaffold3 | 455978 | 456007 | 30 | 2 | 0 | 0 |
| *kl-5* | 2 | Y_scaffold3 | 455825 | 455923 | 99 | 1.92 | 0 | 0 |
| *kl-5* | 3 | Y_scaffold3 | 432310 | 432483 | 174 | 2.42 | 0 | 0 |
| *kl-5* | 4 | Y_scaffold3 | 431995 | 432254 | 260 | 2.15 | 0 | 0 |
| *kl-5* | 5 | Y_scaffold3 | 431598 | 431937 | 340 | 1.86 | 0.04 | 0 |
| *kl-5* | 6 | Y_scaffold3 | 431275 | 431540 | 266 | 1.87 | 0.01 | 0 |
| *kl-5* | 7 | Y_scaffold3 | 430651 | 431218 | 568 | 2.51 | 0.01 | 0.08 |
| *kl-5* | 8 | Y_scaffold3 | 430481 | 430593 | 113 | 3.18 | 0 | 0.91 |
| *kl-5* | 9 | Y_scaffold3 | 430138 | 430429 | 292 | 3.67 | 0.1 | 0.88 |
| *kl-5* | 10 | Y_scaffold3 | 429926 | 430079 | 154 | 4.61 | 0 | 0.91 |
| *kl-5* | 11 | Y_scaffold3 | 351757 | 352587 | 831 | 2.56 | 0 | 0.44 |
| *kl-5* | 12 | Y_scaffold3 | 351249 | 351700 | 452 | 3.8 | 0 | 0.12 |
| *kl-5* | 13 | Y_scaffold3 | 348395 | 351202 | 2808 | 5.25 | 0 | 0.19 |
| *kl-5* | 14 | Y_scaffold3 | 287400 | 289072 | 1673 | 12.38 | 0 | 0.92 |
| *kl-5* | 15 | Y_scaffold3 | 266308 | 266527 | 220 | 12.96 | 0 | 0.66 |
| *kl-5* | 16 | Y_scaffold3 | 254611 | 256649 | 2039 | 14.91 | 0 | 0.91 |
| *kl-5* | 17 | Y_scaffold3 | 201607 | 201745 | 139 | 15.03 | 0 | 0.71 |
| *kl-5* | 18 | Y_scaffold3 | 201382 | 201548 | 167 | 15.68 | 0 | 0.91 |
| *kl-5* | 19 | Y_scaffold3 | 198570 | 201329 | 2760 | 20.22 | 0 | 1.57 |
| *kl-5* | 20 | Y_scaffold3 | 151100 | 151294 | 195 | 100.48 | 0 | 10.62 |
| *kl-5* | 21 | Y_scaffold3 | 51987 | 52086 | 100 | 36.47 | 0 | 3.9 |

Supplemental Table 2. Assembly locations of *PCKR* and *Su(Ste)* island domains and their intra-domain pairwise sequence identity, obtained without gap penalization.

| Strain | Structural_Domain | Copy_Number | Start_Coordinate | End_Coordinate | Intra_Domain_Identity_Pct |
|---|---|---|---|---|---|
| iso-1 | *PCKR_Forward* | 60 | 2852210 | 3103835 | 97.995 |
| iso-1 | *SuSte_1* | 40 | 3103999 | 3290602 | 97.218 |
| iso-1 | *PCKR_Inverted* | 12 | 3292677 | 4344514 | 97.094 |
| iso-1 | *SuSte_2* | 67 | 3363818 | 3555929 | 98.383 |
| iso-1 | *SuSte_3* | 46 | 3675072 | 3889870 | 98.217 |
| iso-1 | *SuSte_4* | 27 | 3937938 | 4012279 | 97.774 |
| iso-1 | *SuSte_5* | 52 | 4118565 | 4330382 | 97.971 |
| iso-1 | *SuSte_6* | 60 | 4365982 | 4865533 | 98.239 |
| iso-1 | *SuSte_7* | 14 | 4865534 | 4930131 | 97.923 |
| iso-1 | *SuSte_8* | 52 | 4943529 | 5460381 | 97.437 |
| A3 | *PCKR_Forward* | 57 | 2006621 | 2228375 | 97.877 |
| A3 | *SuSte_1* | 36 | 2228539 | 2416047 | 97.057 |
| A3 | *PCKR_Inverted* | 17 | 2418122 | 3703794 | 97.177 |
| A3 | *SuSte_2* | 39 | 2497571 | 2616459 | 97.813 |
| A3 | *SuSte_3* | 48 | 2730968 | 2944457 | 98.209 |
| A3 | *SuSte_4* | 72 | 3095619 | 3311399 | 98.31 |
| A3 | *SuSte_5* | 64 | 3417388 | 3689646 | 98.099 |
| A3 | *SuSte_6* | 48 | 3725264 | 4223035 | 97.223 |
| A3 | *SuSte_7* | 15 | 4223036 | 4274416 | 97.948 |
| A3 | *SuSte_8* | 48 | 4287808 | 4796022 | 97.468 |
| A4 | *PCKR_Forward* | 57 | 1975002 | 2212404 | 97.92 |
| A4 | *SuSte_1* | 39 | 2212568 | 2391582 | 97.298 |
| A4 | *PCKR_Inverted* | 15 | 2393658 | 3705548 | 97.284 |
| A4 | *SuSte_2* | 98 | 2450187 | 2745168 | 98.339 |
| A4 | *SuSte_3* | 46 | 2856649 | 3069535 | 98.02 |
| A4 | *SuSte_4* | 73 | 3140348 | 3386027 | 98.141 |

| | | | | | |
|---|---|---|---|---|---|
| A4 | *SuSte_5* | 53 | 3499319 | 3689019 | 97.974 |
| A4 | *SuSte_6* | 70 | 3727024 | 4220450 | 98.063 |
| A4 | *SuSte_7* | 14 | 4220451 | 4278306 | 97.93 |
| A4 | *SuSte_8* | 50 | 4291686 | 4958958 | 97.519 |

Supplemental Table 3. Presence of βNAC*tes1* promoter sequence within each structural island domain across assemblies.

| Strain | Structural_Domain | Total_Units | Units_With_Promoter | Proportion_Pct |
|---|---|---|---|---|
| iso-1 | *PCKR_Forward* | 60 | 0 | 0 |
| iso-1 | *SuSte_1* | 40 | 38 | 95 |
| iso-1 | *PCKR_Inverted* | 12 | 0 | 0 |
| iso-1 | *SuSte_2* | 67 | 2 | 2.99 |
| iso-1 | *SuSte_3* | 46 | 33 | 71.74 |
| iso-1 | *SuSte_4* | 27 | 6 | 22.22 |
| iso-1 | *SuSte_5* | 52 | 36 | 69.23 |
| iso-1 | *SuSte_6* | 60 | 25 | 41.67 |
| iso-1 | *SuSte_7* | 14 | 13 | 92.86 |
| iso-1 | *SuSte_8* | 52 | 50 | 96.15 |
| A3 | *PCKR_Forward* | 57 | 0 | 0 |
| A3 | *SuSte_1* | 36 | 35 | 97.22 |
| A3 | *PCKR_Inverted* | 17 | 0 | 0 |
| A3 | *SuSte_2* | 39 | 2 | 5.13 |
| A3 | *SuSte_3* | 48 | 36 | 75 |
| A3 | *SuSte_4* | 72 | 20 | 27.78 |
| A3 | *SuSte_5* | 64 | 48 | 75 |
| A3 | *SuSte_6* | 48 | 19 | 39.58 |
| A3 | *SuSte_7* | 15 | 14 | 93.33 |
| A3 | *SuSte_8* | 48 | 46 | 95.83 |
| A4 | *PCKR_Forward* | 57 | 0 | 0 |
| A4 | *SuSte_1* | 39 | 38 | 97.44 |
| A4 | *PCKR_Inverted* | 15 | 0 | 0 |

| | | | | |
|---|---|---|---|---|
| A4 | *SuSte_2* | 98 | 2 | 2.04 |
| A4 | *SuSte_3* | 46 | 31 | 67.39 |
| A4 | *SuSte_4* | 73 | 20 | 27.4 |
| A4 | *SuSte_5* | 53 | 36 | 67.92 |
| A4 | *SuSte_6* | 70 | 33 | 47.14 |
| A4 | *SuSte_7* | 14 | 12 | 85.71 |
| A4 | *SuSte_8* | 50 | 48 | 96 |

Supplemental Table 4. Repeat unit size distribution of *PCKR* and *Su(Ste)* repeat units.

| Strain | Type | Size_Category | Count |
|---|---|---|---|
| A3 | *PCKR* | 2300 - 2700 bp | 60 |
| A3 | *PCKR* | < 2300 bp | 9 |
| A3 | *PCKR* | > 2700 bp | 5 |
| A3 | *SuSte* | 2400 - 3000 bp | 304 |
| A3 | *SuSte* | < 2400 bp | 41 |
| A3 | *SuSte* | > 3000 bp | 25 |
| A4 | *PCKR* | 2300 - 2700 bp | 55 |
| A4 | *PCKR* | < 2300 bp | 10 |
| A4 | *PCKR* | > 2700 bp | 7 |
| A4 | *SuSte* | 2400 - 3000 bp | 380 |
| A4 | *SuSte* | < 2400 bp | 39 |
| A4 | *SuSte* | > 3000 bp | 24 |
| iso-1 | *PCKR* | 2300 - 2700 bp | 61 |
| iso-1 | *PCKR* | < 2300 bp | 7 |
| iso-1 | *PCKR* | > 2700 bp | 4 |
| iso-1 | *SuSte* | 2400 - 3000 bp | 299 |
| iso-1 | *SuSte* | < 2400 bp | 35 |
| iso-1 | *SuSte* | > 3000 bp | 24 |

Supplemental Table 5. Location and sequence identity between the inverted *PCKR* units identified as boundaries between *Su(Ste)* island domains. Relative position indicates how far along the array the unit is located, expressed as a percentage of the array.

| Group_ID | Strain | Scaffold | Anchor_Start | Anchor_End | Avg_Identity | Relative_Pos |
|---|---|---|---|---|---|---|
| inv_anchor_group_004 | A3 | Y_scaffold1 | 2497571 | 2500402 | 99.74 | 0.177 |
| inv_anchor_group_004 | A4 | Y_scaffold1 | 2450187 | 2453018 | 99.74 | 0.16 |
| inv_anchor_group_004 | iso-1 | Y_scaffold1 | 3363818 | 3366649 | 99.74 | 0.197 |
| inv_anchor_group_005 | A3 | Y_scaffold1 | 3061230 | 3063650 | 96.59 | 0.379 |
| inv_anchor_group_005 | A4 | Y_scaffold1 | 3105216 | 3107578 | 96.59 | 0.379 |
| inv_anchor_group_005 | iso-1 | Y_scaffold1 | 3907712 | 3910113 | 96.59 | 0.405 |
| inv_anchor_group_006 | A3 | Y_scaffold1 | 3061230 | 3063650 | 97.47 | 0.379 |
| inv_anchor_group_006 | A4 | Y_scaffold1 | 3105216 | 3107578 | 97.47 | 0.379 |
| inv_anchor_group_006 | iso-1 | Y_scaffold1 | 3910113 | 3912533 | 97.47 | 0.406 |
| inv_anchor_group_007 | A3 | Y_scaffold1 | 3691670 | 3694042 | 99.89 | 0.605 |
| inv_anchor_group_007 | A4 | Y_scaffold1 | 3691043 | 3693415 | 99.89 | 0.575 |
| inv_anchor_group_007 | iso-1 | Y_scaffold1 | 4332406 | 4334778 | 99.89 | 0.568 |
| inv_anchor_group_009 | A3 | Y_scaffold1 | 3691670 | 3694042 | 99.61 | 0.605 |
| inv_anchor_group_009 | A4 | Y_scaffold1 | 3691043 | 3693415 | 99.61 | 0.575 |
| inv_anchor_group_009 | iso-1 | Y_scaffold1 | 4337254 | 4339613 | 99.61 | 0.57 |
| inv_anchor_group_012 | A3 | Y_scaffold1 | 4287808 | 4290668 | 95.55 | 0.818 |

| | | | | | | |
|---|---|---|---|---|---|---|
| inv_anchor_group_012 | A4 | Y_scaffold1 | 4368698 | 4371503 | 95.55 | 0.803 |
| inv_anchor_group_012 | iso-1 | Y_scaffold1 | 4943529 | 4946389 | 95.55 | 0.802 |
| inv_anchor_group_013 | A3 | Y_scaffold1 | 4287808 | 4290668 | 95.39 | 0.818 |
| inv_anchor_group_013 | A4 | Y_scaffold1 | 4371504 | 4374320 | 95.39 | 0.804 |
| inv_anchor_group_013 | iso-1 | Y_scaffold1 | 4946390 | 4949244 | 95.39 | 0.803 |
| inv_anchor_group_014 | A3 | Y_scaffold1 | 4287808 | 4290668 | 95.04 | 0.818 |
| inv_anchor_group_014 | A4 | Y_scaffold1 | 4374321 | 4377126 | 95.04 | 0.805 |
| inv_anchor_group_014 | iso-1 | Y_scaffold1 | 4949245 | 4952109 | 95.04 | 0.805 |
| inv_anchor_group_027 | A3 | Y_scaffold1 | 4338810 | 4341655 | 97.26 | 0.837 |
| inv_anchor_group_027 | A4 | Y_scaffold1 | 4460727 | 4463554 | 97.26 | 0.834 |
| inv_anchor_group_027 | iso-1 | Y_scaffold1 | 5025010 | 5027862 | 97.26 | 0.834 |
| inv_anchor_group_029 | A3 | Y_scaffold1 | 4338810 | 4341655 | 95.76 | 0.837 |
| inv_anchor_group_029 | A4 | Y_scaffold1 | 4468542 | 4471380 | 95.76 | 0.836 |
| inv_anchor_group_029 | iso-1 | Y_scaffold1 | 5032584 | 5035400 | 95.76 | 0.837 |
| inv_anchor_group_059 | A3 | Y_scaffold1 | 4481349 | 4483989 | 97.3 | 0.888 |
| inv_anchor_group_059 | A4 | Y_scaffold1 | 4502767 | 4505505 | 97.3 | 0.848 |
| inv_anchor_group_059 | iso-1 | Y_scaffold1 | 5165156 | 5167796 | 97.3 | 0.887 |
| inv_anchor_group_060 | A3 | Y_scaffold1 | 4487068 | 4489701 | 96.11 | 0.89 |
| inv_anchor_group_060 | A4 | Y_scaffold1 | 4502767 | 4505505 | 96.11 | 0.848 |
| inv_anchor_grou | iso-1 | Y_scaffol | 5170874 | 5173507 | 96.11 | 0.89 |

| | | | | | | |
|---|---|---|---|---|---|---|
| p_060 | | d1 | | | | |
| inv_anchor_group_064 | A3 | Y_scaffold1 | 4493165 | 4495541 | 99.97 | 0.999 |
| inv_anchor_group_064 | A4 | Y_scaffold1 | 4500390 | 4502766 | 99.97 | 0.999 |
| inv_anchor_group_064 | iso-1 | Y_scaffold1 | 5175263 | 5177639 | 99.97 | 0.999 |
| inv_anchor_group_065 | A3 | Y_scaffold1 | 4495542 | 4498280 | 99.85 | 1 |
| inv_anchor_group_065 | A4 | Y_scaffold1 | 4502767 | 4505505 | 99.85 | 1 |
| inv_anchor_group_065 | iso-1 | Y_scaffold1 | 5177640 | 5180375 | 99.85 | 1 |

Supplemental Table 6. Copy number variation between X-linked *Stellate* Euchromatic and Heterochromatic units between iso-1, A3, and A4 strains, annotated according to Shukla et al., 2025. Highlights sequence identity between units within each respective domain via all-versus-all pairwise alignments.

| Strain | *Stellate* variant | Copy Number | Mean_Internal_Identity_Pct |
|---|---|---|---|
| iso-1 | *HetSte_L1* | 1 | 100 |
| iso-1 | *EuSte_Distal* | 1 | 100 |
| iso-1 | *EuSte_Proximal* | 10 | 94.031 |
| iso-1 | *HetSte_Singleton* | 1 | 100 |
| iso-1 | *HetSte_L2* | 4 | 98.463 |
| iso-1 | *HetSte_L3* | 11 | 99.096 |
| iso-1 | *HetSte_L4* | 1 | 100 |
| iso-1 | *HetSte_L5* | 1 | 100 |
| iso-1 | *HetSte_L6* | 1 | 100 |
| iso-1 | *HetSte_L7* | 1 | 100 |
| A3 | *EuSte_Distal* | 1 | 100 |
| A3 | *EuSte_Proximal* | 1 | 100 |
| A3 | *HetSte_L1* | 1 | 100 |

| | | | |
|---|---|---|---|
| A3 | *HetSte_L2* | 4 | 98.463 |
| A3 | *HetSte_L3* | 11 | 99.045 |
| A3 | *HetSte_L4* | 1 | 100 |
| A3 | *HetSte_L5* | 1 | 100 |
| A3 | *HetSte_L6* | 1 | 100 |
| A3 | *HetSte_L7* | 1 | 100 |
| A4 | *EuSte_Distal* | 1 | 100 |
| A4 | *EuSte_Proximal_Main* | 188 | 98.475 |
| A4 | *EuSte_Proximal_Island* | 9 | 91.343 |
| A4 | *HetSte_Singleton_1* | 1 | 100 |
| A4 | *HetSte_Singleton_2* | 1 | 100 |

Supplemental Table 7. All-versus-all pairwise sequence identity between X and Y-linked rDNA repeat units in each assembly, categorized by the presence of R1 and R2 TE insertions. Canonical units are units without transposon insertions in the coding sequence backbone.

| **iso-1** | Column 1 | Column 2 | Column 3 | Column 4 | Column 5 |
|---|---|---|---|---|---|
| Chromosome | rDNA variants | Pair Count | Average Identity | Median Identity | Range (Min-Max) |
| Intra-Y | Canonical vs Canonical | 231 | 99.80% | 99.87% | 98.96% – 100% |
| | R1 vs R1 | 136 | 99.66% | 99.75% | 99.21% – 100% |
| | R2 vs R2 | 378 | 91.35% | 99.65% | 43.78% – 100% |
| | R1 vs Canonical | 374 | 99.69% | 99.76% | 98.56% – 100% |
| | R2 vs Canonical | 616 | 95.70% | 99.78% | 51.67% – 99.99% |
| | R1 vs R2 | 476 | 92.83% | 97.19% | 49.52% – 99.95% |
| Intra-X | Canonical vs Canonical | 3 | 90.51% | 87.30% | 84.91% – 99.32% |
| | R1 vs R1 | 136 | 98.64% | 99.82% | 88.64% – 99.98% |
| | R2 vs R2 | 3 | 99.31% | 99.03% | 99.02% – |

| | | | | | |
|---|---|---|---|---|---|
| | | | | | 99.87% |
| | R1 vs Canonical | 51 | 92.02% | 99.13% | 73.02% – 99.86% |
| | R2 vs Canonical | 9 | 93.91% | 99.44% | 81.61% – 99.69% |
| | R1 vs R2 | 51 | 93.13% | 93.54% | 82.14% – 99.81% |
| Inter-X/Y | Canonical vs Canonical | 66 | 95.37% | 99.46% | 86.54% – 99.84% |
| | R1 vs R1 | 289 | 98.98% | 99.55% | 88.61% – 99.84% |
| | R2 vs R2 | 84 | 95.01% | 99.20% | 50.92% – 99.89% |
| | R1 vs Canonical | 425 | 98.24% | 99.46% | 77.63% – 99.82% |
| | R2 vs Canonical | 150 | 93.81% | 99.50% | 46.07% – 99.86% |
| | R1 vs R2 | 527 | 88.48% | 89.00% | 45.55% – 99.81% |
| **A3** | | | | | |
| Section | Comparison Pair | Pair Count | Average Identity | Median Identity | Range (Min-Max) |
| Intra-Y | Canonical vs Canonical | 820 | 93.18% | 99.85% | 49.19% – 100% |
| | R1 vs R1 | 66 | 99.62% | 99.69% | 98.82% – 99.96% |
| | R2 vs R2 | 190 | 94.67% | 99.54% | 51.71% – 100% |
| | R1 vs Canonical | 492 | 95.77% | 99.63% | 51.19% – 100% |
| | R2 vs Canonical | 820 | 93.57% | 99.55% | 48.13% – 100% |
| | R1 vs R2 | 240 | 91.94% | 95.45% | 54.47% – 99.96% |
| Intra-X | Canonical vs Canonical | 45 | 99.81% | 99.88% | 99.45% – 100% |
| | R1 vs R1 | 406 | 90.41% | 99.61% | 50.93% – 100% |
| | R2 vs R2 | 55 | 83.63% | 99.37% | 50.32% – 100% |
| | R1 vs Canonical | 290 | 94.76% | 99.67% | 51.75% – 100% |
| | R2 vs Canonical | 110 | 91.08% | 99.81% | 51.72% – 100% |

| | | | | | |
|---|---|---|---|---|---|
| | R1 vs R2 | 319 | 84.55% | 92.81% | 49.19% – 99.98% |
| Inter-X/Y | Canonical vs Canonical | 410 | 96.12% | 99.61% | 51.74% – 99.92% |
| | R1 vs R1 | 348 | 94.56% | 99.55% | 50.93% – 99.92% |
| | R2 vs R2 | 220 | 88.93% | 99.48% | 49.77% – 100% |
| | R1 vs Canonical | 1309 | 92.26% | 99.48% | 49.13% – 99.97% |
| | R2 vs Canonical | 651 | 90.98% | 99.47% | 48.06% – 99.97% |
| | R1 vs R2 | 712 | 87.96% | 91.98% | 49.10% – 100% |
| **A4** | | | | | |
| Section | Comparison Pair | Pair Count | Average Identity | Median Identity | Range (Min-Max) |
| Intra-Y | Canonical vs Canonical | 6 | 99.61% | 99.78% | 99.10% – 99.96% |
| | R1 vs R1 | 3 | 99.83% | 99.77% | 99.77% – 99.96% |
| | R2 vs R2 | 21 | 99.42% | 99.50% | 98.32% – 99.94% |
| | R1 vs Canonical | 12 | 99.63% | 99.77% | 98.84% – 99.97% |
| | R2 vs Canonical | 28 | 99.42% | 99.71% | 97.40% – 100% |
| | R1 vs R2 | 21 | 89.89% | 88.82% | 86.69% – 94.46% |
| Intra-X | Canonical vs Canonical | 15 | 99.63% | 99.77% | 99.17% – 99.92% |
| | R1 vs R1 | 66 | 90.59% | 98.46% | 51.55% – 99.95% |
| | R2 vs R2 | 21 | 99.74% | 99.74% | 99.57% – 99.91% |
| | R1 vs Canonical | 72 | 94.88% | 99.75% | 52.50% – 99.95% |
| | R2 vs Canonical | 42 | 99.67% | 99.68% | 99.38% – 99.98% |
| | R1 vs R2 | 84 | 87.80% | 89.52% | 52.22% – 98.32% |

| | | | | | |
|---|---|---|---|---|---|
| Inter-X/Y | Canonical vs Canonical | 24 | 99.45% | 99.77% | 97.97% – 99.88% |
| | R1 vs R1 | 36 | 95.37% | 99.73% | 52.51% – 99.81% |
| | R2 vs R2 | 49 | 99.52% | 99.62% | 97.81% – 99.90% |
| | R1 vs Canonical | 66 | 96.20% | 99.67% | 52.46% – 99.88% |
| | R2 vs Canonical | 70 | 99.45% | 99.60% | 97.65% – 99.89% |
| | R1 vs R2 | 105 | 87.41% | 88.87% | 48.07% – 97.06% |

Chang C-H, Larracuente AM. 2019. Heterochromatin-Enriched Assemblies Reveal the Sequence and Organization of the Drosophila melanogaster Y Chromosome. *Genetics* 211:333–348.

Schauer T, Ghavi-Helm Y, Sexton T, Albig C, Regnard C, Cavalli G, Furlong EE, Becker PB. 2017. Chromosome topology guides the Drosophila Dosage Compensation Complex for target gene activation. *EMBO Rep.* 18:1854–1868.

Shukla HG, Chakraborty M, Emerson JJ. 2025. Genetic variation in recalcitrant repetitive regions of the Drosophila melanogaster genome. *Genome Res.* 35:2023–2040.
